# IGHMBP2 deletion suppresses translation and activates the integrated stress response

**DOI:** 10.1101/2023.12.11.571166

**Authors:** Jesslyn E. Park, Hetvee Desai, José Liboy-Lugo, Sohyun Gu, Ziad Jowhar, Albert Xu, Stephen N. Floor

## Abstract

IGHMBP2 is a non-essential, superfamily 1 DNA/RNA helicase that is mutated in patients with rare neuromuscular diseases SMARD1 and CMT2S. IGHMBP2 is implicated in translational and transcriptional regulation via biochemical association with ribosomal proteins, pre-rRNA processing factors, and tRNA-related species. To uncover the cellular consequences of perturbing IGHMBP2, we generated full and partial IGHMBP2 deletion K562 cell lines. Using polysome profiling and a nascent protein synthesis assay, we found that IGHMBP2 deletion modestly reduces global translation. We performed Ribo-seq and RNA-seq and identified diverse gene expression changes due to IGHMBP2 deletion, including ATF4 upregulation. With recent studies showing the ISR can contribute to tRNA metabolism-linked neuropathies, we asked whether perturbing IGHMBP2 promotes ISR activation. We generated ATF4 reporter cell lines and found IGHMBP2 knockout cells demonstrate basal, chronic ISR activation. Our work expands upon the impact of IGHMBP2 in translation and elucidates molecular mechanisms that may link mutant IGHMBP2 to severe clinical phenotypes.

## INTRODUCTION

IGHMBP2 is a ubiquitously-expressed, superfamily 1 (SF1) DNA/RNA helicase implicated in the regulation of mRNA translation (Grohmann et al. 2001; de Planell-Saguer et al. 2009). IGHMBP2 mutations cause the rare autosomal recessive diseases spinal muscular atrophy with respiratory distress type 1 (SMARD1) and Charcot-Marie-Tooth disease type 2S (CMT2S), which are characterized by severe neurodegeneration and myopathies (Grohmann et al. 2001; Cottenie et al. 2014). The disease-associated nonsense, frameshift, or missense mutations in IGHMBP2 are predicted to impair DNA/ RNA binding, unwinding, and ATP hydrolysis, and thus pathogenicity is attributed to loss-of-function of IGHMBP2 enzymatic activity (Guenther et al. 2009a; Lim et al. 2012; Saladini et al. 2020). While the mechanism of cellular pathogenesis upon functional loss of IGHMBP2 is unknown, IGHMBP2-dependent translational disruption is a favorable hypothesis considering the relevance of translational dysregulation among inherited disease mechanisms (Scheper et al. 2007). For example, a prevalent cause of amyotrophic lateral sclerosis (ALS) are expansions in *C9orf72,* which lead to global translational repression that inhibits the mRNA decay activity of UPF1, another SF1 helicase (Sun et al. 2020). Defective helicases cause numerous translation-linked pathologies beyond neurodegeneration—DDX3X is a primary example of an SF2 helicase associated with neurodevelopmental disorders and cancers when its translational activities are perturbed (Bohnsack et al. 2023; Gadek et al. 2023).

To-date, the connection between IGHMBP2 and translational regulation has been evidenced by prior biochemical work and using model mice or mouse motoneurons. IGHMBP2 is predominantly cytoplasmic, where it co-localizes with translation initiation factor eIF4G2 and rRNA (Grohmann et al. 2004; Guenther et al. 2009a). Therefore, despite IGHMBP2 having *in vitro* activity on DNA or RNA, its physiological substrate may be primarily RNA. IGHMBP2 co-precipitates and co-sediments with ribosomal proteins and subunits, and thus interacts with translational machinery during translation (Guenther et al. 2009a). Localized translation, as indicated by expression of a translationally-regulated β-actin GFP-3′UTR reporter, is slowed in IGHMBP2-deficient motoneurons post-photobleaching (Surrey et al. 2018). IGHMBP2 also co-precipitates with macromolecules that contribute to translational regulation such as elongation factors and tRNA-Tyr (Guenther et al. 2009a; de Planell-Saguer et al. 2009). Mice with spontaneous IGHMBP2 mutations–which model human SMARD1 phenotypes–were found to be rescued when expressing a genetic modifier locus encoding ABT1 and several tRNA genes, further implicating a connection between tRNAs and IGHMBP2 in disease pathology (Cox et al. 1998; de Planell-Saguer et al. 2009).

Recent work has established a connection between tRNA and diverse neurodegenerative disorders. Disruption in tRNA regulatory genes via mutations in various aminoacyl-tRNA synthetases (ARS) causes several axonal types of CMT, such as CMT2N, CMT2U, CMT2W, and CMT2D, and can also cause other forms of neurodegeneration or ataxia (Lee et al. 2006). Recently, defective GARS activity in CMT2D was found to induce ribosomal stalling and activate GCN2, triggering the integrated stress response (ISR) (Spaulding et al. 2021; Mendonsa et al. 2021). The ISR mediates cellular reprogramming toward pro-survival or apoptosis in response to various exogenous or endogenous stimuli, including translation elongation stalls due to tRNA deficiency, and is a disease mechanism among several neurodegenerative disorders (Costa-Mattioli & Walter, 2020). It is unclear whether the ISR is activated by loss of IGHMBP2.

In this study, we investigated the physiological and molecular consequences of IGHMBP2 deletion in human cells. We found that IGHMBP2 depletion causes cell proliferation and global protein synthesis to decline, supporting a role of IGHMBP2 in global translational efficiency. We defined the gene expression changes caused by loss of IGHMBP2 using RNA-seq and ribosome profiling, and found that expression of the stress-induced transcription factor ATF4 was upregulated in IGHMBP2 deletion cells. We generated ATF4 reporter cell lines to confirm the occurrence and reversibility of chronic, IGHMBP2-associated activation of the ISR. Our observation of ISR activation and translational suppression caused by IGHMBP2 deletion may point to a possible disease mechanism and solidifies a role for IGHMBP2 in translational regulation.

## RESULTS

### IGHMBP2 deletion impacts cellular proliferation and global translation

To investigate the molecular function of IGHMBP2, we deleted IGHMBP2 in K562 CRISPRi cells using CRISPR-Cas9 with two sgRNAs targeting exon 2 of *IGHMBP2* (Fig. 1A). Edited clones were screened for inactivating alleles via PCR, and selected partial and full-knockout IGHMBP2 monoclonal cell lines were validated by Sanger sequencing and Western blot (Fig. 1B,C; Supplemental Fig. S1A-E). To determine the physiological impact of IGHMBP2 deletion, we stably expressed mEGFP in IGHMBP2 deletion clones and measured competitive proliferation relative to parental cells via flow cytometry across two weeks of cell passages. We observed decreased relative cell size and lower proliferation rates among IGHMBP2 full-deletion cells (Fig. 1D,E; Supplemental Fig. S1F,G). We confirmed mEGFP expression does not impact cell proliferation and that cell lines expressing an alternate fluorophore expressed in subsequent experiments, TagBFP, exhibited comparable proliferation results (Supplemental Fig. S1H). Expression of each fluorescent protein remained robust across the experimental timeframes (Supplemental Fig. S1I). We conclude that IGHMBP2 loss induces a modest proliferation defect in K562 cells.

**Figure 1.**
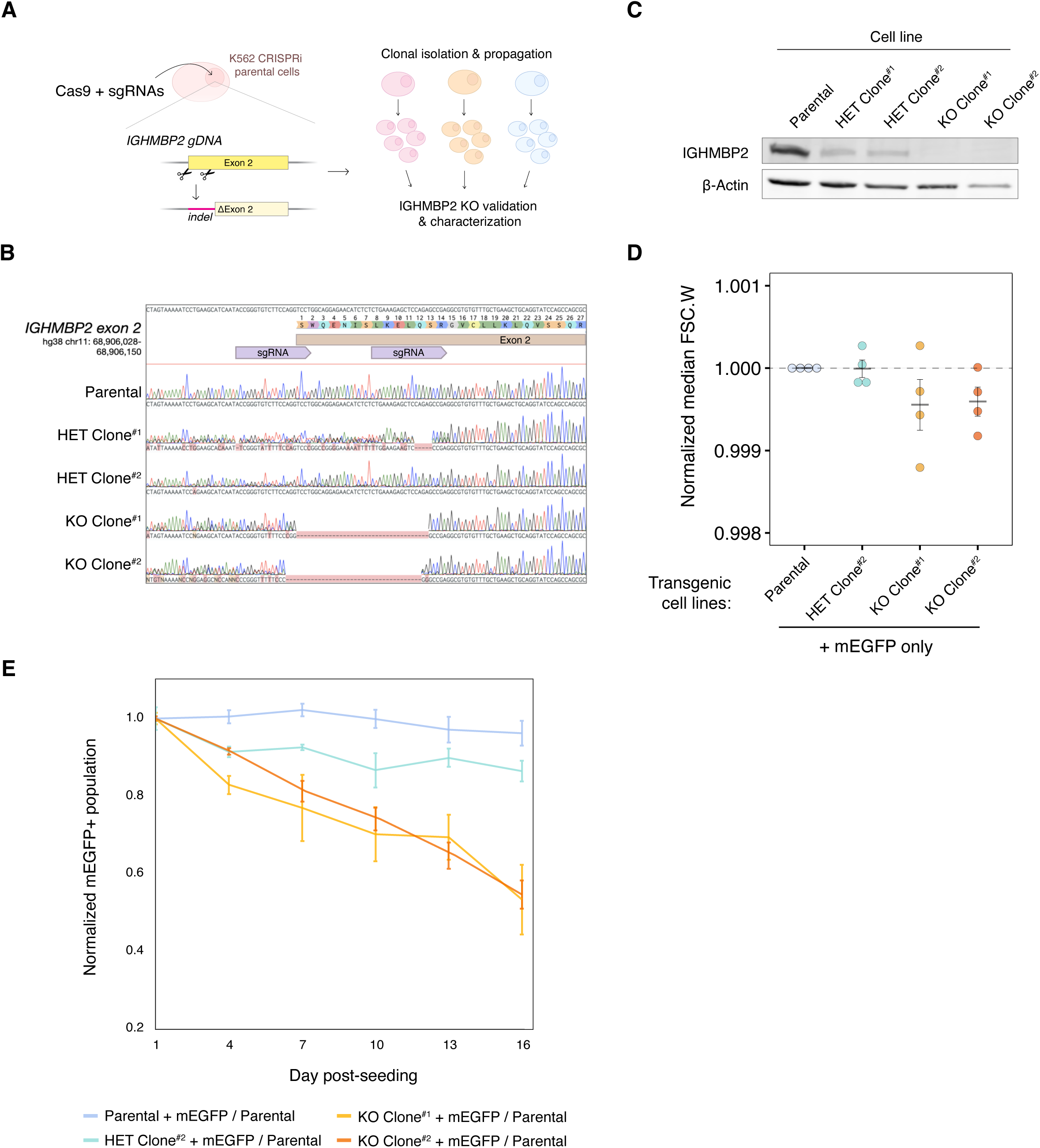
IGHMBP2 deletion decreases proliferation of cells. (*A*) Cas9-mediated IGHMBP2 deletion in K562 CRISPRi cells. (*B*) Sanger sequencing of gDNA surrounding IGHMBP2 exon 2 cut-site in clones screened for indels via PCR. sgRNA guide sequences used to dually target Cas9 are indicated in purple. (*C*) Western blot of IGHMBP2 expression in clones depicted in B, validating reduced IGHMBP2 protein levels in HET Clones #1 and #2 and full deletion in KO Clones #1 and #2. (*D*) Forward-scatter width medians between cell lines with differential IGHMBP2 expression normalized to the median FSC.W of parental cells per day of measurement. (*E*) Competitive proliferation profiles between L’IGHMBP2 cell lines stably expressing mEGFP seeded with 50% non-fluorescent parental cells. Each sample was seeded in triplicate on Day 0 and independently passaged on each day of measurement. Error bars reflect the coefficient of variation mEGFP+ populations normalized to the mean Day 1 reading among triplicate wells per sample.

The effect of IGHMBP2 on proliferation could be related to its described role in protein synthesis. We therefore assessed the impact of IGHMBP2 deletion on global translation by performing polysome profiling (Fig. 2A). Western blot of polysome fractions shows IGHMBP2 co-sediments with 40S, 60S, and 80S ribosomal species (Fig. 2B). When we quantified the area under the curve (AUC) per polysome and compared relative peak ratios per cell line, we observed mild yet reproducible enrichment of free ribosomal subunits relative to monosomes and polysomes in IGHMBP2 deletion clones, while subunit-specific abundances held constant. This suggests translational suppression caused by IGHMBP2 deletion is not due to gross ribosomal biogenesis defects (Fig. 2C). To monitor global translation rates in cell lines differentially expressing IGHMBP2 at single-cell resolution, we measured relative rates of nascent polypeptide synthesis via an O-propargyl-puromycin (OPP) assay. We found translation was suppressed in IGHMBP2 deletion clones (Fig. 2D; Supplemental Fig. S2), consistent with the polysome profiling results. We reproduced these results in cell lines expressing transgenic mEGFP, which were initially sorted to include all GFP+ cells, and detected decreased mEGFP expression per cell in IGHMBP2-disrupted clones as an additional orthogonal measure of translational activity (Fig. 2 E,F).

**Figure 2.**
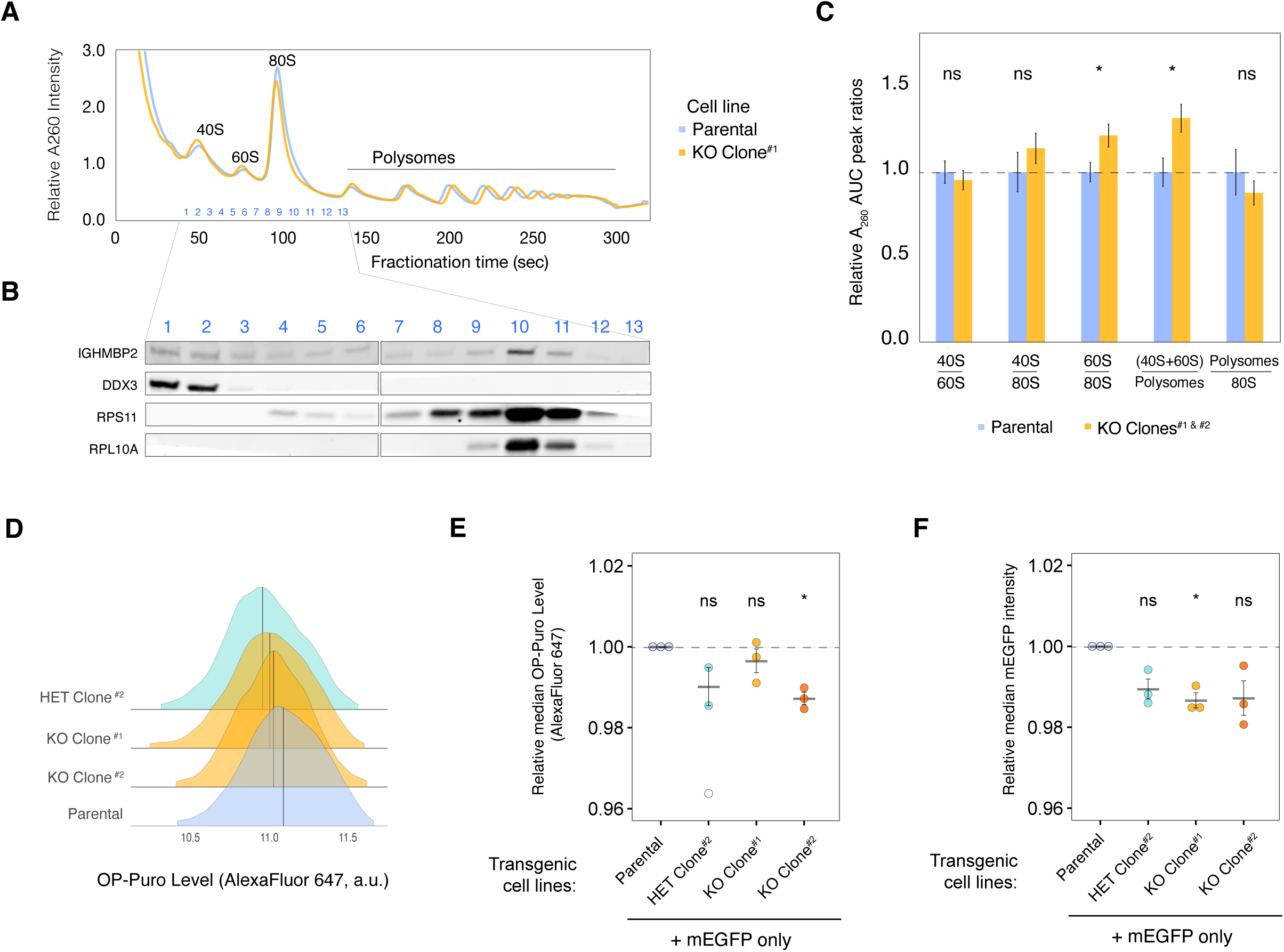
IGHMBP2 deletion reduces global translation in cells. (*A*) Representative polysome profiles from parental K562 CRISPRi (blue) and IGHMBP2 KO Clone (yellow). (*B*) Western blot of fractions of the polysome profile in A. (*C*) Area under the curve (AUC) quantification of polysome profiles, where n = 5 for parental samples and n = 7 for IGHMBP2 deletion clone samples collected from separate flasks and measured across 3 different days. Unpaired, two-tailed, two-sample t-test between parental and KO samples per AUC ratios shown was performed to determine significance, and error bars represent standard error of the mean (SEM). (*D*) Representative OP-Puro (OPP) levels of cell lines used in this study quantified via flow cytometry, which was performed in 3 independent experiments with either AlexaFluor 647 as shown or AlexaFluor 594 (*Fig. S2*) (*E*) OPP assay results using cell lines expressing mEGFP only. Values beyond 2.5 standard deviations (SD) from the mean of all data points were considered outliers omitted in statistical measurement (SD = 2.6 for single outlier in HET clone; unfilled). (*F*) Relative mEGFP expression in samples from *E*. For *E* and *F*, n = 3, where n represents median fluorescence of single-cell intensities normalized to the median fluorescence of parental cells from experiments on 3 separate days. Horizontal black lines per each sample represent the mean, and error bars are SEM. Median values were derived from at least 10,000 cells measured by flow cytometry. Unpaired t-test was performed to determine significance. For *B* and *F,* ns: p ≥; 0.05, *: p ≤ 0.05.

### Ribosome profiling reveals differentially expressed genes in IGHMBP2 deletion cells

Modest global changes in translation can result from changes to the translation of a subset of mRNAs (Calviello & Venkataramanan et al. 2021). To determine the gene expression changes in IGHMBP2 deletion cells, we performed ribosome profiling alongside RNA-seq. Ribo-seq and RNA-seq read counts showed high correlation between replicates (Supplemental Fig. S3). We confirmed Ribo-seq data exhibited 3-nt periodicity, and the majority of reads were forward-sense stranded and mapped to coding exons as expected (Supplemental Figs. S4-S6). When broadly comparing changes between datasets, we observed an increased magnitude and number of gene expression changes in full versus partial IGHMBP2 deletion cells (Fig. 3A; Supplemental Table 1). To interpret the mode of gene regulation as a consequence of IGHMBP2 deletion, we classified genes according to adjusted p-values derived from differential expression (DE) analysis. For RNA-seq and Ribo-seq results, log_2_ foldchanges (L2FC) were determined per gene relative to parental expression levels, and genes with IGHMBP2-dependent RNA:Ribo-seq interaction term changes were classified under Δtranslational efficiency (ΔTE). To gain further insight into the expression changes underlying ΔTE, ΔTE genes were categorized as translation exclusive, where only ribosome occupancy level was changed significantly per gene, or translation buffered, where only differential abundance of mRNA was observed (Chothani et al. 2019; Supplemental Table 1).

**Figure 3.**
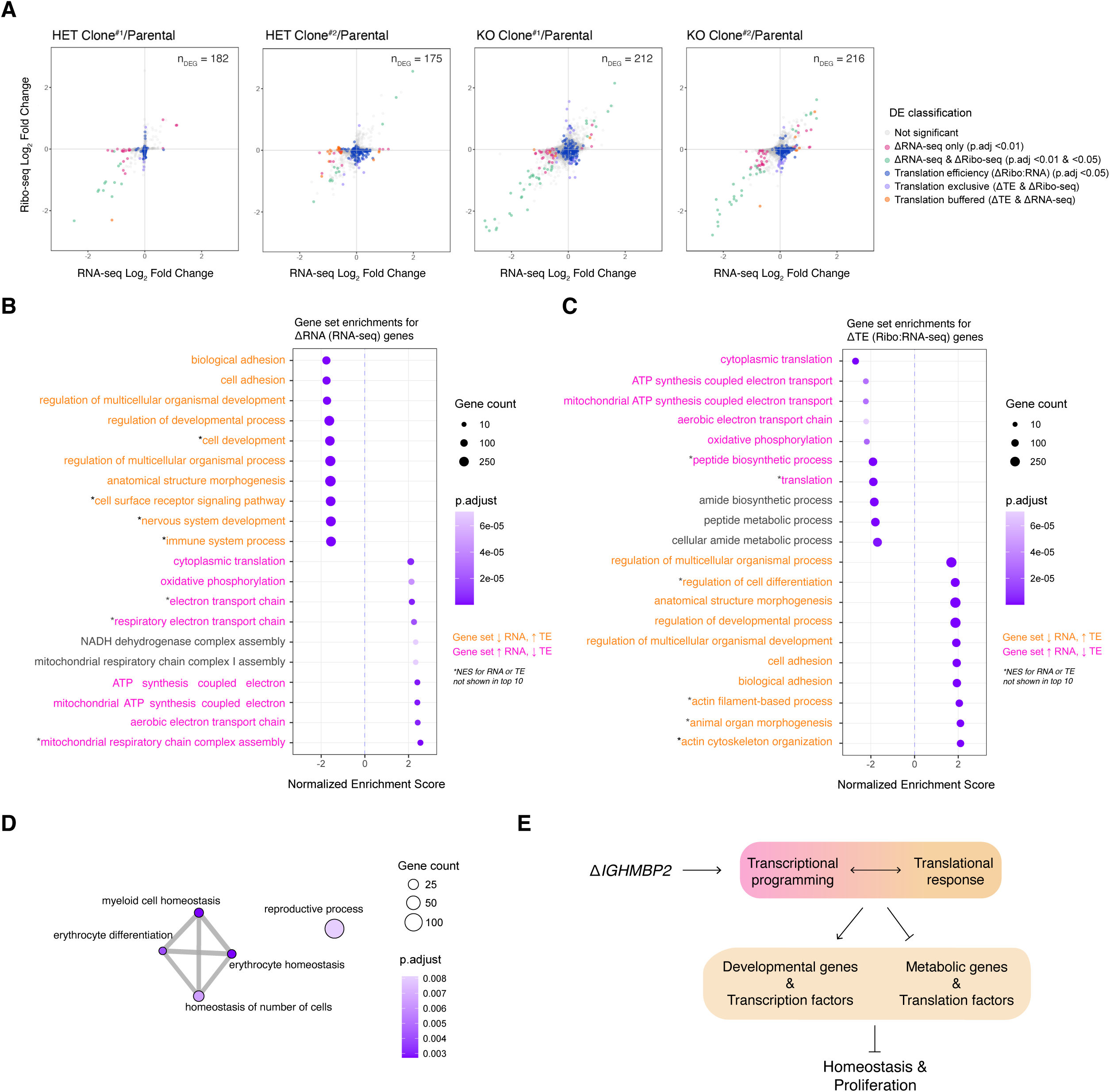
IGHMBP2 loss alters translation of diverse mRNAs. (A) Ribosome profiling versus RNA-seq-derived shrunken log_2_ (Fold Change) (L2FC) per gene in clones with partial or full IGHMBP2 deletion compared to parental cells. Differential expression (DE) analysis was performed using Wald test, and p-values were adjusted (p.adj) via Benjamini-Hochberg method. Cut-offs used for DE classifications are p.adj <0.01 for LiRNA-seq (pink, green, and orange), and p.adj <0.05 for LiRibo-seq (green & violet) and Litranslation efficiency (LiTE; blue, violet, and orange). Genes with LiTE across all clones were identified via likelihood ratio test against the Ribo:RNA-seq interaction term across all samples. Genes of both LiTE and LiRibo-seq are identified as translation exclusive (violet). Genes of Li TE and LiRNA-seq are classified as translation buffered (orange). The number of DEGs (n_DEG_) are shown per cell line. (B) Top 10 up versus down-regulated gene set enrichments (GSE) using ranked DEGs from RNA-seq and (*C*) Ribo:RNA-seq. GS labels are pink or orange if expression trends between *B* and *C* are opposing positively or negatively, respectively. (*D*) Enrichment network map of GSEA with Ribo-seq reads. In *B*, *C*, and *D*, GSEA was computed using the Biological Processes ontology; min GS size = 25, max GS size = 1,000 with 100,000 permutations. (*E*) Overview of physiological impact of IGHMBP2 disruption.

We functionally characterized genes differentially expressed among IGHMBP2 deletion clones via gene set enrichment analysis (GSEA) using fold-change data from RNA-seq and Ribo-seq DE results (Supplemental Table 2). We observed gene expression attenuation between ΔmRNA and ΔTE gene sets: translation machinery and mitochondrial gene sets were transcriptionally upregulated in IGHMBP2 deletion cells, yet suppressed in ΔTE (Fig. 3B). Developmental and homeostatic gene sets were transcriptionally suppressed, yet increased in ΔTE in IGHMBP2 deletion cells (Fig. 3C). We note that 167 out of 500 possible genes in the peptide biosynthetic process, which largely overlaps with translation gene sets, were suppressed in IGHMBP2 KO clones, comprising ribosomal machinery, translation initiation, and translation elongation factors. The regulation of developmental processes gene set was highly upregulated in TE due to IGHMBP2 deletion, comprising 306 of 854 possible genes, which broadly overlapped with upregulated genes in the anatomical structure morphogenesis gene set. Beyond sets comprising diverse signaling genes, the negative regulation of DNA-binding transcription factor gene set was also significantly upregulated with a normalized enrichment score (NES) of 1.6, reflecting 236 of 582 possible genes (Supplemental Table 2). Upon performing GSEA with ribosomal footprint fold-changes, we observed translational suppression of various homeostatic gene sets appropriately corresponding with the hemopoietic cell type of K562 cells (Fig. 3D). Together, these data identify the genetic changes underlying morphological and translation defects observed as a consequence of IGHMBP2 deletion, again implicating translation as a process impacted by loss of IGHMBP2 (Fig. 3E).

### ATF4 is upregulated in IGHMBP2 deletion cells

Considering the recessive nature of IGHMBP2 pathology, we sought to determine the genetic changes specific to full loss of IGHMBP2 to identify possible disease-relevant targets. Of several ΔTE genes identified from our RNA:Ribo-seq analysis (Supplemental Fig. S7A), ATF4 expression was notably upregulated in genes filtered for dependence on full loss of IGHMBP2 (Fig. 4A). Upon assessing TE classification, ATF4 was classified as translation exclusive ΔTE gene only in the full IGHMBP2 KO genotype condition (Supplemental Fig. S7B-C). Induction of ATF4 is caused by translational control through two upstream open reading frames (uORF; Lu et al. 2004; Vattem & Wek 2004). We observed an enrichment of normalized reads mapped to uORF1 and uORF2 of *ATF4* in IGHMBP2 deletion clones relative to parental cells, especially at the enhancer uORF1 (Fig. 4B,C), suggesting translational activation of ATF4 expression.

**Figure 4.**
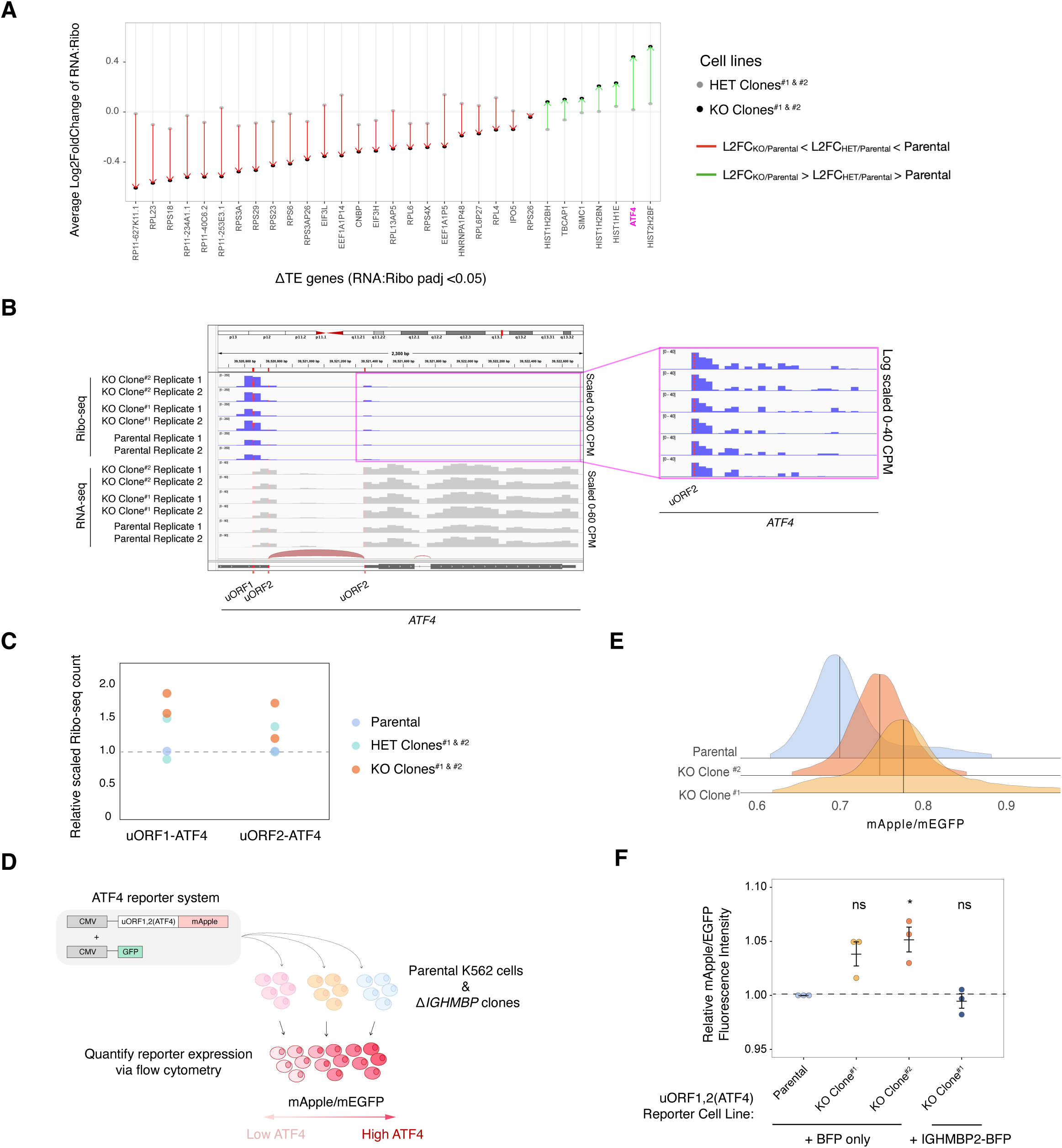
ATF4 is upregulated in IGHMBP2 deletion cells. (*A*) Average L2FC of a subset of /1TE DE genes in heterozygous and full deletion clones relative to parental cells. DE genes were filtered to represent those of <10% FC in heterozygous IGHMBP2 cells, where L2FC in HET clones were intermediate relative to KO results. (*B*) Mapped reads normalized to counts per million surrounding the 5’-UTR of ATF4. (*C*) Quantification of reads aligning to either of the two uORFs in ATF4 in the parental, HET clones, and KO clones. Each dot represents the average scaled count per clones between two replicates. (*D*) Schematic of ATF4 reporter system. Transcription of lentivirally-integrated mApple and GFP reporter constructs is driven by separate CMV promoters. An ISR-sensitive, synthetic 5’-UTR encoding two uORFs (derived from ATF4) is upstream of the mApple ORF. (*E*) uORF1,2(ATF4)-mApple expression normalized to promoter and translational activity (mEGFP) in /1IGHMBP2 K562 reporter cell lines. (*F*) Relative median mApple/mEGFP intensities among reporter cell lines expressing BFP or IGHMBP2-BFP. Single-cell mApple/mEGFP signals were normalized to the median mApple/mEGFP of parental cells from 3 different days (n = 3). Median values were derived from at least 10,000 cells measured by flow cytometry. Unpaired t-test was performed to determine significance (ns: p ≥ 0.05, *: p ≤ 0.05, **: p ≤ 0.01).

To test the hypothesis that IGHMBP2 deletion is inducing ATF4 expression, we generated parental and ΔIGHMBP2 reporter cell lines stably expressing a CMV-driven (uORF1,2)-ATF4-mApple reporter gene (Fig. 4D). As part of the reporter system, a separate CMV-GFP transgene was introduced to enable normalization for promoter strength and translational activity (Guo et al. 2020). We used this reporter due to challenges with robust detection of ATF4 via immunoblot. We detected elevated ATF4 reporter levels in IGHMBP2 deletion clones relative to parental (Fig. 4E; Supplemental Fig. S8A,B). We confirmed inducibility of reporter expression upon acute ISR stimulation via thapsigargin (Supplemental Fig. S8C). We then exogenously expressed IGHMBP2 N- or C-terminally fused with TagBFP in our IGHMBP2 deletion reporter cell lines and observed ATF4 reporter expression reversed toward parental levels of ATF4 with both transgenes (Fig. 4F; Supplemental Fig. S8D,E), indicating genetic rescue and supporting the causative nature of *IGHMBP2* genetic lesions. We therefore find that loss of IGHMBP2 induces ATF4 expression.

### IGHMBP2 deletion results in ISR activation

ATF4 is a stress-responsive transcriptional regulator that is tightly regulated at the translational level (Harding et al. 2000; Vattem & Wek 2000). Given the role of ATF4 as part of the ISR and the relevance of the ISR in neurodegenerative diseases, we wondered if activation of the ISR is a consequence of IGHMBP2 deletion. Surprisingly, differences in p-eIF2α were not discernible between parental and IGHMBP2 deletion cells via Western blot (Supplemental Fig. S9A-B). This may be due to homeostatic regulation, which produces reduced levels of p-eIF2α upon chronic ISR activation relative to acute stress (Novoa et al. 2003; Guan et al. 2017). However, upon treating IGHMBP2 deletion cell lines with p-eIF2α-dependent ISR inhibitor ISRIB, ATF4 reporter expression in IGHMBP2 KO cells robustly reversed toward parental reporter levels at steady-state, supporting the occurrence of chronic, p-eIF2α-induced ISR activation (Fig. 5A).

**Figure 5.**
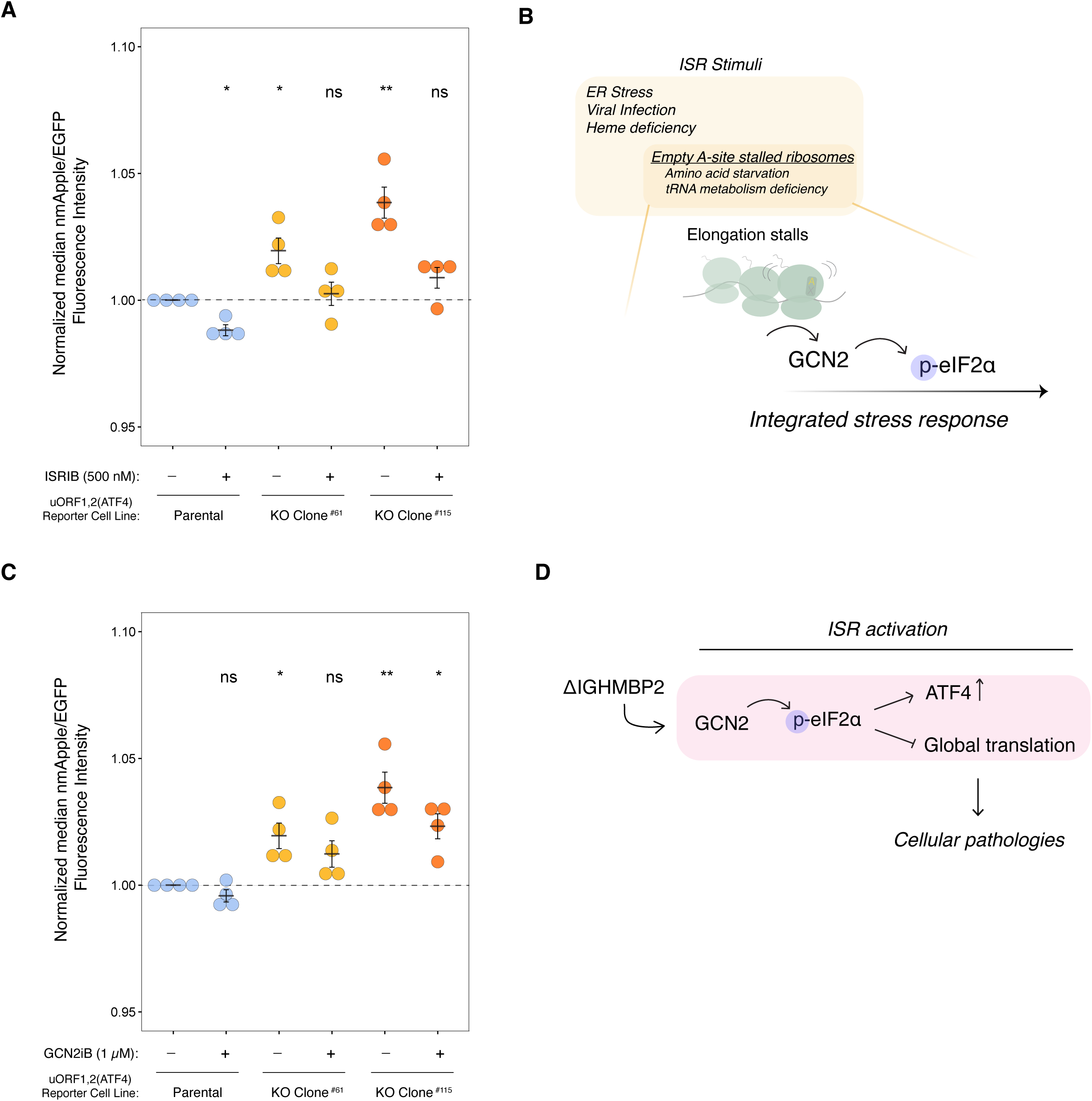
The ISR is activated in IGHMBP2 deletion cells. (*A*) Relative median mApple/mEGFP fluorescence intensities among reporter cell lines treated with 500 nM ISRIB for 24 hr, where single-cell mApple/mEGFP signals were normalized to the median mApple/mEGFP of untreated (DMSO only) parental cells. (*B*) Schematic of GCN2-mediated ISR activation. (*C*) Relative median mApple/mEGFP fluorescence intensities among reporter cell lines treated with 1 µM GCN2iB for 24 hr, normalized as described in *A*. For *A* and *D*, n=4 where n reflects results from separate experiments performed on different days, and error bars represent SEM. Unpaired t-test was performed to determine significance (ns: p ≥ 0.05, *: p ≤ 0.05, **: p ≤ 0.01). (*D*) A model of how IGHMBP2 deletion results in chronic ISR activation.

Four kinases can activate the ISR: PKR senses specific types of dsRNA, HRI senses mitochondrial dysfunction, PERK responds to the unfolded protein response in the endoplasmic reticulum, and GCN2 activates upon ribosome collisions induced by starvation or deficiencies in tRNA metabolism (Costa-Mattioli & Walter, 2020). Given the genetic connection between IGHMBP2 and tRNA metabolism, we hypothesized ISR activation in IGHMBP2 deletion cells could be driven by empty A-site induced ribosomal stalling and downstream GCN2 activation (Fig. 5B). We thus treated ATF4 reporter cell lines with kinase inhibitor GCN2iB (Nakamura et al. 2018) and observed a modest decrease in ATF4 reporter expression in IGHMBP2 deletion cell lines (Fig. 5C). Upon determining that ISR activation is driven by GCN2 in IGHMBP2 deletion cells, we wondered if we could detect ribosome stalling among our Ribo-seq reads, as empty A-site stalled and collided ribosomes canonically activate GCN2 (Ishimura et al. 2016; Inglis et al. 2019). We first performed a metagene analysis using Ribo-seq reads to determine whether IGHMBP2 deletion alters ribosome positioning along transcripts globally, as might occur under strong stalling conditions. We observed consistent ribosomal occupancy across sequenced transcripts in knockout compared to parental cell lines (Supplemental Fig. S10A), suggesting IGHMBP2 deletion may impact translation by a more ubiquitous mechanism non-specific to a particular stage of translation, or perhaps toward specific transcripts undetectable by global analysis. Determining whether IGHMBP2-RNA recognition or unwinding activity impacts global translation and how this may be governed by particular biophysical properties or motifs among mRNAs warrants further experimentation.

As tRNA dysregulation may be a favorable hypothesis driving global translation changes, specifically tRNA-Tyr as implicated from prior literature (de Planell-Saguer et al. 2009), we performed a Northern blot to visualize total tRNA-Tyr and tRNA-Val expression among our cell lines (Supplemental Fig. S10B). We did not discern differences in abundance of these tRNAs, suggesting IGHMBP2-dependent differences in tRNA-Tyr abundance may not occur in K562 cells or may occur below detectability among biochemical assays. To further test the hypothesis that IGHMBP2 loss affects decoding of a subset of codons, we applied a recently-developed machine learning package, *choros,* to infer in-frame ribosome A-site occupancy of codons globally, but did not detect specific codon enrichment (Mok et al. 2023; Supplemental Fig. S10C-H). This lack of signal is perhaps due to the subtle effect size of IGHMBP2 loss on translation, or could occur for other unknown reasons. We conclude IGHMBP2 deletion results in chronic, low-level ISR activation mediated by GCN2 signaling (Fig. 5D), which could reflect low-level ribosome collisions below our detection threshold.

## DISCUSSION

In this work, we defined the impact of IGHMBP2 in translation using Cas9-mediated IGHMBP2 knockout human cell lines. We also studied the effects of complete loss of IGHMBP2 expression compared to heterozygous IGHMBP2 cell lines to reflect a biologically-relevant, non-pathological regime. We show that relative to partial deletion or transgenic knock-in, full loss of IGHMBP2 slows cell proliferation, reduces translation, disrupts gene expression across the transcriptome and translatome, and activates the ISR.

To further characterize ISR activation in IGHMBP2 deletion cells, we performed GSEA with a custom gene set comprising 122 compiled ATF4 target genes (Neill & Masson, 2023). While we observed enrichment in ΔTE among ATF4 target genes to a similar magnitude of top gene sets from the Biological Processes gene ontology, this result did not reach statistical significance (Supplemental Fig. S11A,B). ISR-induced apoptotic signaling typically involves CHOP upregulation (Gachon et al. 2001), which we did not detect in our DEG dataset, suggesting cellular stress caused by IGHMBP2 deletion induces a pro-survival ISR program. We notably observed chronic, low-level activation of ATF4 among our IGHMBP2 deletion cell lines, which is also associated with pro-survival ISR programming (Pakos-Zebrucka et al. 2016). We surmise IGHMBP2 deletion-induced, pro-survival ISR may enable cells to respond to upstream molecular consequences of IGHMBP2 disruption, ultimately resulting in subtle rather than severe fitness defects.

Although IGHMBP2 is ubiquitously expressed, IGHMBP2-associated pathogenesis is thought to be tissue-specific. IGHMBP2 is most highly expressed in the brain, and pathological symptoms of SMARD1 and CMT2S are restricted to neural and muscle tissue (Cox et al. 1998; Guenther et al. 2009b; Cottenie et al. 2014). We speculate this may explain the small effect sizes observed in our model in contrast to more measurable ISR effects observed in neuronal cells modeling CMT2D (Spaulding et al. 2021; Zuko et al. 2021). Alternatively, the ISR may be more strongly activated by IGHMBP2 loss in highly proliferative cells such as K562s, thus measuring ISR activation in animal models will be important work for the future. Despite the small effect sizes, we robustly and reproducibly observe modest translation repression and chronic ISR activation using high-throughput and sensitive approaches such as deep sequencing and flow cytometry. Our work may also inform future experimental considerations toward detecting disease-relevant, chronic stress markers that may otherwise remain undetected using traditional techniques.

While tRNA metabolism-driven ISR activation is an intriguing disease mechanism hypothesis, axonal CMT is notably caused by various other sources of molecular dysfunction as well. For example, disruption in mitochondrial activity (CMT2A, CMT2DD, CMT2EE), cytoskeletal proteins (CMT2B1, CMT2E, CMT2CC), and other diverse cellular functionalities converge among CMT pathogenesis. Thus, it is feasible that alternative consequences of IGHMBP2 deletion beyond tRNA dysregulation may more dominantly contribute to neuropathy and myopathy, though it is appealing to speculate this may be tissue-dependent considering SMARD1 model mice are rescued when expressing a modifier locus encoding ABT1 and five tRNA-Tyr genes (de Planell-Saguer et al. 2009). How mechanisms of IGHMBP2-associated pathogenesis may compound or differentially manifest across different tissue types will be important to deconvolve in future studies.

Consequences of IGHMBP2-DNA impairment remain unaddressed in our study and would be important to distinguish considering IGHMBP2 is also found in the nucleus (Grohmann et al. 2004). The namesake of IGHMBP2 refers to its recognition of DNA motifs that aid immunoglobulin heavy chain recombination, and IGHMBP2 exhibits slower, short-lived dsDNA unwinding processivity *in vitro* relative to the prototypical SF1 helicase, UPF1 (Mizuta et al. 1993; Kanaan et al. 2018). Expanded functional insight toward IGHMBP2-DNA and its unique processive behavior is yet to be determined. Whether differential transcription caused by IGHMBP2 deletion precedes cellular stress activation and translational dysregulation remains unclear, and this is confounded by transcriptional reprogramming chronically mediated by the ISR. Future DNA-binding experiments may elucidate the possible relevance of IGHMBP2-DNA disruption toward cellular pathology.

One of 95 broadly elusive, non-redundant human helicases, IGHMBP2 is an understudied non-essential gene with recent findings that shed light on its fundamental importance (Umate et al. 2011). IGHMPB2 was predicted to contribute to cellular fitness in a CRISPR screen in HEK293T cells (Hart et al. 2015), and we establish that perturbing IGHMBP2 indeed also reduces cellular fitness in K562 cells via translation suppression and ISR activation. Beyond the physiological consequences we have identified, a recent screen performed in HeLa cells showed IGHMBP2 disruption alters nucleolar dynamics, further demonstrating its ubiquitous impact across non-neuronal cellular models (Sheu-Gruttadauria et al. 2023). This study also predicted gene functionality based on how genetic perturbations influenced nucleolar dynamics, which categorized IGHMBP2 in a non-ribosomal biogenesis role that disrupts nucleolar rRNA composition, in-line as a potential consequence of IGHMBP2-associated global translation suppression. Promisingly, adeno-associated virus (AAV) therapeutic strategies have been shown to rescue SMA-related phenotypes, including SMARD1 in mice using AAV9-IGHMBP2 (Pattali et al. 2019; Shababi et al. 2016). Thus, our demonstration of reversible IGHMBP2-dependent pathologies may present a therapeutically-relevant, cell-based system for further study. Defining the fundamental function of IGHMBP2 and repercussions immediately downstream of IGHMBP2 deletion will also be imperative for identifying alternative, drug-based therapeutics. Our work demonstrates the utility of studying neuropathy-associated gene IGHMBP2 in an immature human cell line, expands on the fundamental relevance of IGHMBP2, and reinforces the significance of translation in cellular health.

## MATERIALS & METHODS

### Cell culture

K562 CRISPRi cell lines used in this study were maintained in RPMI 1640 (Gibco) containing L-glutamine and 25 mM HEPES, supplemented with 10% FBS and 1% penicillin-streptomycin. Cells were stored in humidified incubators at 37 °C with 5% CO_2_ and routinely checked for mycoplasma (Lonza).

### Cell line generation

Cas9-mediated knockout of IGHMBP2 was performed using two sgRNAs (Synthego) to target exon 2 of IGHMBP2 in combination: 5′-CAGAGAGAUGUUCUCCUGCC-3′ and 5′-CUGAAAGAGCUCCAGAGCCG-3′. Ribonucleoprotein (RNP) complexes were prepared by combining 50 pmol of recombinant Cas9 to 50 pmol of each sgRNA and incubating the mixture for 20 min at 37 °C. K562 CRISPRi cells were nucleofected (SF Cell Line 4D kit; Lonza) with the RNP. Nucleofected clones were isolated by limiting dilution and indel-containing clonal populations were screened by extracting gDNA using Quick Extract (Biosearch Technologies) for PCR amplification using primers targeting up and downstream the Cas9 cut-site: 5′-GGTTGT GGCATTAACTGCCC-3′ and 5′-CCCACATCAATTGTTGGAC-3′. PCR products were separated on a 3% agarose gel at 100 V, and *IGHMBP2* gDNA disruption in select clones was further validated via Sanger sequencing with primer 5′-CTTTAC GAGGGTACAAGTCACGG-3′ and Western blotting.

### Western blot

Cell lines were harvested during active growth (0.4E6 - 0.6E6 cells/mL) and lysed in RIPA buffer supplemented with protease inhibitor cocktail (Halt). Lysates were centrifuged at 16,000 *xg* for 20 min at 4 °C, and total protein concentration was quantified by Pierce BCA Assay (Thermo). SDS loading buffer was added to 50 µg total protein, and samples were boiled at 95 °C for 5 min. Samples were loaded on 4-15% pre-cast PAGE gels (Bio-Rad), run at 85 V in running buffer, and transferred to a PVDF membrane (Bio-Rad). Membranes were incubated for 1 hour in blocking buffer (1% (w/ v) BSA in 0.1% TBS-T), followed by an overnight incubation at 4 °C in blocking buffer containing primary antibody. Membranes were washed for 10 min 0.1% TBS-T three times, followed by a 1 hr incubation with secondary antibody with rotation at room temperature and protected from light. Membranes were washed for 10 min 0.1% TBS-T three times and imaged using an LI-COR Odyssey DLx.

Antibodies used for Western blotting include rabbit anti-IGHMBP2 at 1:500 (*Proteintech*, 23945-1-AP), rabbit anti-β-actin conjugated to AlexaFluor680 (Abcam); rabbit anti-eIF2α at 1:1,000; rabbit anti-eIF2α phospho-S51 at 1:1,000. Membranes were incubated with goat anti-rabbit 800 CW secondary antibody (*LI-COR*, 926-32211) at 1:10,000.

### Northern blot

Northern blotting was performed using NorthernMax kit (Invitrogen) with adaptations as described. Total RNAs were extracted from K562 parental, KO clone #1 and #2 cells with Trizol reagent (Invitrogen,15596026). 5 µg of total RNA per sample was mixed with 100 mM final concentration of Tris-HCl (pH 9.0) and incubated for 30 min at 37 °C to remove amino acids attached to the 3′ end of tRNAs. Samples were then mixed with 2X Formaldehyde Load Dye (Invitrogen, 8550G) and heat denatured for 3 min at 70 °C. RNAs were loaded onto pre-cast 10% TBE-Urea gels (Thermo Fisher, EC68755BOX) in 1X TBE. RNAs were separated by PAGE, transferred to a positively charged nylon membrane (Invitrogen, AM10104) and cross-linked to the membrane using a UV trans-illuminator (Stratalinker 2400 UV Crosslinker) for 2 min at 1200 x 100 µJ. Membranes were pre-hybridized with pre-warmed ULTRAhybTM ultrasensitive hybridization buffer (Invitrogen, AM8670) at 52 °C for 1 hr, followed by addition of 10 pmol of each probe. The membranes were hybridized overnight and washed two times with Low Stringency Wash buffer (Invitrogen, AM8673) with rotation at room temperature for 5 min. Membranes were probed with either tRNA-Tyr, tRNA-Val, or tRNA-Val, stripped, re-probed for tRNA-Gly as an internal control, and imaged using an LI-COR Odyssey DLx.

tRNA-Tyr and tRNA-Val probes were labeled with IRDye800RD and the tRNA-Gly probe was labeled with IRDye680RD (LI-COR), respectively. Probe sequences used include Gly-sCC (5′-TCATTGGCCRGGAATYGAACCCGGGYCTCCCRCGT GGWAGGCGAGAATTCTACCACTGMACCACCMAYG/iAzideN/C-3′); Tyr-GTA: (5′-TCC TTCGAGCCGGASTCGAACCAGCGACCTAAGGATCTACAGTCCTCCGCTCTACCARCTGAGCTATCGAAG/iAzideN/G-3′); Val-mAC: (5′-TGTTTCYGCCYGGTTTCG AACCRGGGACCTTTCGCGTGTKAGGCGAACGTGATAACCACTACACTACRGAAA/ iAzideN/C-3 ′); Val-TAC (5 ′-TGGTTCCACTGGGGCTCGAACCCAGGACCTTCTGCGTGTAAAGCAGACGTGATAACCACTACACTATGGAAC/iAzideN/C-3′) (Dittmar et al. 2006).

### Transgenic cell line generation

For virus generation, Lenti-X HEK 293T cells (Takara Bio) were transfected with lentiviral packaging plasmids and transgenic constructs. Virus was harvested and clarified through a 0.45 µm filter after 3 days. Lentiviral titering was optimized to result in <50% transduction to generate cells with MOI <1, and cells were infected via centrifugation at 150 *xg* for 1 hr in 8 µg/mL polybrene (EMD Millipore). After 3 days, transduced cells were selected via FACS using a 85-100 µm diameter nozzle.

### Flow cytometry

Flow cytometry was performed on BD Fortessa instruments. When the high-throughput screening (HTS) 96-well plate attachment was utilized, plate parameters were set to a flow rate of 0.5-1.0 µL/sec, 20 µL sample volume, 50 µL mixing volume, 200 µL/sec mixing speed, 2 mixes, and 200 µL wash volume. Laser parameters were adjusted to approximately 400 V for FSC, 250 V for SSC, 260 V for Violet laser detected with 450/50 bandpass filter (for BFP), 350 V for Blue detected with 530/30 bandpass filter (for mEGFP and AlexaFlour488), 580 V for YGD detected with 586/615 bandpass filter (for mApple), 520 V for YGC detected with 610/620 bandpass filter (for AlexaFlour594), 320 V for Red detected with 670/630 bandpass filter (for AlexaFlour647).

### Nascent protein synthesis assay

The Click-iT Plus OPP Protein Synthesis Assay Kit (*ThermoFisher*, C10457) was utilized as described in a 96-well format with 0.5E6 cells harvested during active growth per reaction. Cells were co-incubated with OPP in DMSO or 125 nM cycloheximide as a negative control for 30 minutes prior to fixation with 4% PFA and permeabilization with 0.25% Triton-X 100 in PBS. Cells were resuspended in PBS with 5% (w/v) FBS and differential fluorescent signal between cell lines was observed by flow cytometry.

### Cell proliferation competition assay

Cell lines were seeded in 96-well plates at 1.5E5 cells/mL. In wells where transgenic cell lines were seeded with parental cells, 50% abundance of non-parental cells was targeted. Flow cytometry data was collected one day after seeding, showing 35-50% starting populations of fluorescent cells compared to parental cells. Wells were passaged for subsequent measurements every 3 days for 16 days.

### Polysome profiling

Cells were grown in separate flasks in triplicate and treated with 100 μg/mL cycloheximide for 5 min. 10E6 cells were harvested per sample and lysed with a hypotonic lysis buffer (10 mM HEPES pH 7.9, 1.5 mM MgCl_2_, 10 mM KCl, 0.5 mM DTT, 1% Triton X-100, 100 µg/mL cycloheximide) and trituration with a 26 G needle. Sucrose gradients were prepared with 10% and 50% sucrose in sucrose gradient buffer (100 mM KCl, 20 mM HEPES pH 7.6, 5 mM MgCl2, 1 mM DTT, 100 µg/mL cycloheximide). For each cell line, 100 µL lysate was layered atop the sucrose gradient, and polysomal species were separated by ultracentrifugation at 36,000 RPM for 2 hours at 4 °C. Polysome profiles were obtained via injection through a spectrophotometer at a flow rate of 2 mL/min, and absorbance was recorded at 260 nm with sensitivity set to 0.1 on the UV/Vis detector.

### Ribosome profiling

Cells were seeded in fresh media 16 hr prior to harvest. Cell count was measured near 0.3E6 cells/mL per cell line at time of harvest, reflecting active growth conditions. Ribosomal footprint RNAs were obtained as previously described (Calviello & Venkataramanan et al. 2021). Cell lysates was treated with RNase I and subjected to size exclusion chromatography using MicroSpin Columns S-400 HR (Illustra). Ribosome-protected RNA was then extracted with Trizol, and RNA corresponding to the size of monosomal footprints (26-34 nt) were isolated by size selection via running RNA samples in a 15% polyacrylamide TBE-Urea gel. Footprint fragments were dephosphorylated and ligated to pre-adenylated oligonucleotide linker (NI-816: 5′-/ 5Phos/NNNNNTAGACAGATCGGAAGA GCACACGTCTGAA/3ddC/-3′) with Mth RNA ligase. Unligated linker was removed with RecJ exonuclease. Reverse transcription was performed on linker-ligated RNA with Protoscript II (primer NI-802: 5′-/5Phos/ NNAGATCGGAAGAGCGTCGTGTAGGGAAAGAG/iSp18/GTGA CTGGAGTTCAGACG TGTGCTC-3′), and samples were treated with 1 M NaOH to hydrolyze remaining RNAs. cDNA was size-selected with a 15% polyacrylamide TBE-Urea gel and circularized with CircLigase II. rRNA was depleted from the sample with a subtraction oligo pool, and cDNA was quantified via qPCR. cDNA libraries were amplified with Phusion polymerase (Forward primerNI-798: 5 ′-AATGATACGGCGACCACCGAGATCTACACTCTTTCCCTACACGACGCTC) using unique reverse index primers (Reverse primer: 5′- CAAGCAGAAGACGGCATACGAGATJJJJJJGTG ACTGGAGTTCAGACGTGTG-3′) and size-selected with a 15% polyacrylamide TBE-Urea gel. The quality and concentration of amplified libraries was assessed with the Bioanalyzer and Qubit; 5 ng/sample were pooled and sequenced with the Illumina HiSeq 4000 (SE65 66×8×8×0) via the sequencing core of the UCSF Center for Advanced Technology.

### RNA-seq

Total RNA was extracted from lysates using Direct-zol RNA Miniprep kit (Zymo) and NEBNext Ultra II Directional RNA Illumina Library Prep Kit (NEB E7760) with NEBNext rRNA Depletion Kit V2 (NEB E7400). RNA-seq samples were prepared with single index primers (NEB ME6609S). RNA integrity and and concentration of amplified libraries was assessed with the Bioanalyzer and Qubit; 5 ng/sample were pooled and sequenced with the Illumina HiSeq 4000 (SE65 66×8×8×0) via the sequencing core of the UCSF Center for Advanced Technology.

### Sequencing pre-processing

FASTQ files were converted to FASTA format. Adapter sequences from ribo-seq FASTQ files were removed with *cutadapt*. Using *cutadapt*, adapter sequences were removed from RNA-seq and Ribo-seq sample reads, and reads were filtered to a minimum length of 22 nt. Reads were collapsed by UMI, which were then removed. Ribosomal RNAs, repeat RNAs and other non-coding RNAs were aligned against and removed with *repeatmasker* using bowtie2 2.4.1. Index files were generated using STAR 2.7.5a, GRCh38 primary assembly genome.fa file, and a GENCODE v25 .gtf annotation file with --sjdbOverhang set to 64 or 29 for RNA-seq and Ribo-seq indices, respectively. Filtered reads were then mapped to corresponding indices, generating annotated .bam files for analysis.

### Sequencing analysis

RNA-seq and ribo-seq results were analyzed using *DESeq2* library in R (Love et al., 2014). Transcript biotypes from the corresponding GTF annotation (GENCODE v25) were matched to all genes, and “rRNA” and “Mt_rRNA”-classified genes remaining after pre-processing were removed. Mapped reads were pre-filtered by performing DESeq analysis on all samples to determine the minimal filterThreshold. The minimal filterThreshold was determined to be 3.7 for RNA-seq samples and 5.8 for Ribo-seq samples, and thus genes with counts below these cut-offs were removed from corresponding datasets. DESeq was run with pre-filtered genes with independentFiltering omitted, as this widened ability to statistically contextualize DE changes among genes otherwise abundantly filtered out with high independent filterThresholds in nuanced perturbation conditions, especially among heterozygous clones. Dispersion results were then shrunken using the *apeglm* method.

Because we compare more than two groups (parental, KO, and HET genotypes) a likelihood ratio test (LRT) was employed rather than default Wald testing, which uses standard error. The initial design used was ∼ SeqType + Condition + SeqType:Condition, and DESeq was run using likelihood ratio testing (LRT) with a reduced design ∼SeqType + Condition to derive DE results reflecting SeqType:Condition (Ribo:RNA with respect to genotype). DESeq was run with filterThreshold pre-filtered genes with independentFiltering omitted. Due to the modeling design, resultant p-values are the same between KO or HET clones, and L2FC were independently determined among samples. Results were then filtered by baseMean cut-offs of >50 for RNA-seq and baseMean >40 for Ribo-seq. Genes were classified as significant for RNA-seq p-adjusted values <0.01 and Ribo-seq p-adjusted values <0.05. To further account for possible clonal artifacts and outliers, final significant DE genes among RNA-seq and Ribo-seq results must be significant in at least 2 out of 4 unique clones; genes found significant in only 1 out of the 4 clonal cell lines were filtered from the final datasets.

R packages used to assess quality control of sequencing data include *MultiQC*, *RiboprofilingQC*, and *RiboseQC*. To determine localized read mapping coverage for specific genes, such as *ATF4* uORFs, bamCoverage from *deeptools* was used to convert RNA-seq and Ribo-seq pre-processed *.bam* files to *.bigwig* files, where reads were normalized to counts per million (CPM). CPM-scaled reads were then mapped to hg38 using Interactive Genomics Viewer (IGV).

### Data availability

Sequencing data is available at GEO accession number GSE248890.

### Gene set enrichment analysis

Using RNA-seq, Ribo:RNA-seq, and Ribo-seq DEG lists output from DESEQ analysis, the L2FC per gene was averaged between cell lines. Mean L2FC were sorted as a ranked list, and GSEA was performed using *fgsea* and *clusterprofiler* R packages using the Biological Process sub-ontology. The minimum and maximum geneset sizes were set to 25 and 1,000, respectively. GSEA was performed with 100,000 permutations, and exported genesets reflect a p-value cutoff of 0.01. The enrichment map plot was generated using the *enrichplot* library.

### Metagene analysis

Metagene analysis was performed using *ribosomeProfilingQC* library in R with .bam files and the coverageDepth function. The GENCODE v25 .gtf annotation file was used to define UTR and CDS regions. Metagene profiles were then Min-Max normalized between 0 and 1 for comparability between each clone.

### Codon enrichment analysis

Ribo-seq reads were mapped to the transcriptome using STAR 2.7.5a -- quantMode TranscriptomeSAM. Transcriptomic .bam files were assigned posterior probability values using RSEM (rsem-calculate-expression) with --fragment-length-mean 29 and then input into the *choros* R library. For parental and KO Clone #2 samples, A-site offset rule values were determined to be 15, 14, and 16 for frame 0, 1, and 2 for reads between lengths 29-30 nt; 15, 14, and NA for frame 0, 1, and 2 for 28 nt reads; NA, 14, and NA for frame 0, 1, and 2 for 27 nt reads, respectively. A-site offset rules were less discernible in KO Clone #1, and thus was omitted from codon enrichment analysis.

## ACKNOWLEDGEMENTS

Sequencing was performed by the UCSF CAT, supported by UCSF PBBR, RRP IMIA, and NIH 1S10OD028511-01 grants. Flow cytometry and cell sorting was performed at the UCSF Parnassus Flow Cytometry CoLab. K-562 CRISPRi cells were gifted by the Jonathan Weissman lab. We thank Matthew Taliaferro and Wilfried Rossoll for crucial conversations during the formation of this project. The authors also thank the Floor lab members for support and helpful feedback, especially Jess Sheu-Gruttadauria, Yizhu Lin, and Sam Kwok for technical advice. This work was supported by the National Institutes of Health R35GM149255 (to S.N.F.). S.N.F. is a Pew Scholar in the Biomedical Sciences, supported by The Pew Charitable Trusts.

## Author contributions

J.E.P generated cell lines, performed experiments, and conducted data analysis. H.D. supported initial knockout cell line characterization, Western blotting, and polysome profiling. J.L. supported sequencing pre-processing. S.G. performed Northern blotting. Z.J. and A.X. shared computational expertise and provided technical support toward cloning and cell culture. All authors contributed thoughtful discussion. J.E.P and S.N.F co-conceptualized experiments and co-wrote the manuscript, which was revised by all authors. S.N.F. acquired funding and supervised the project.

---

**Figure S1.**
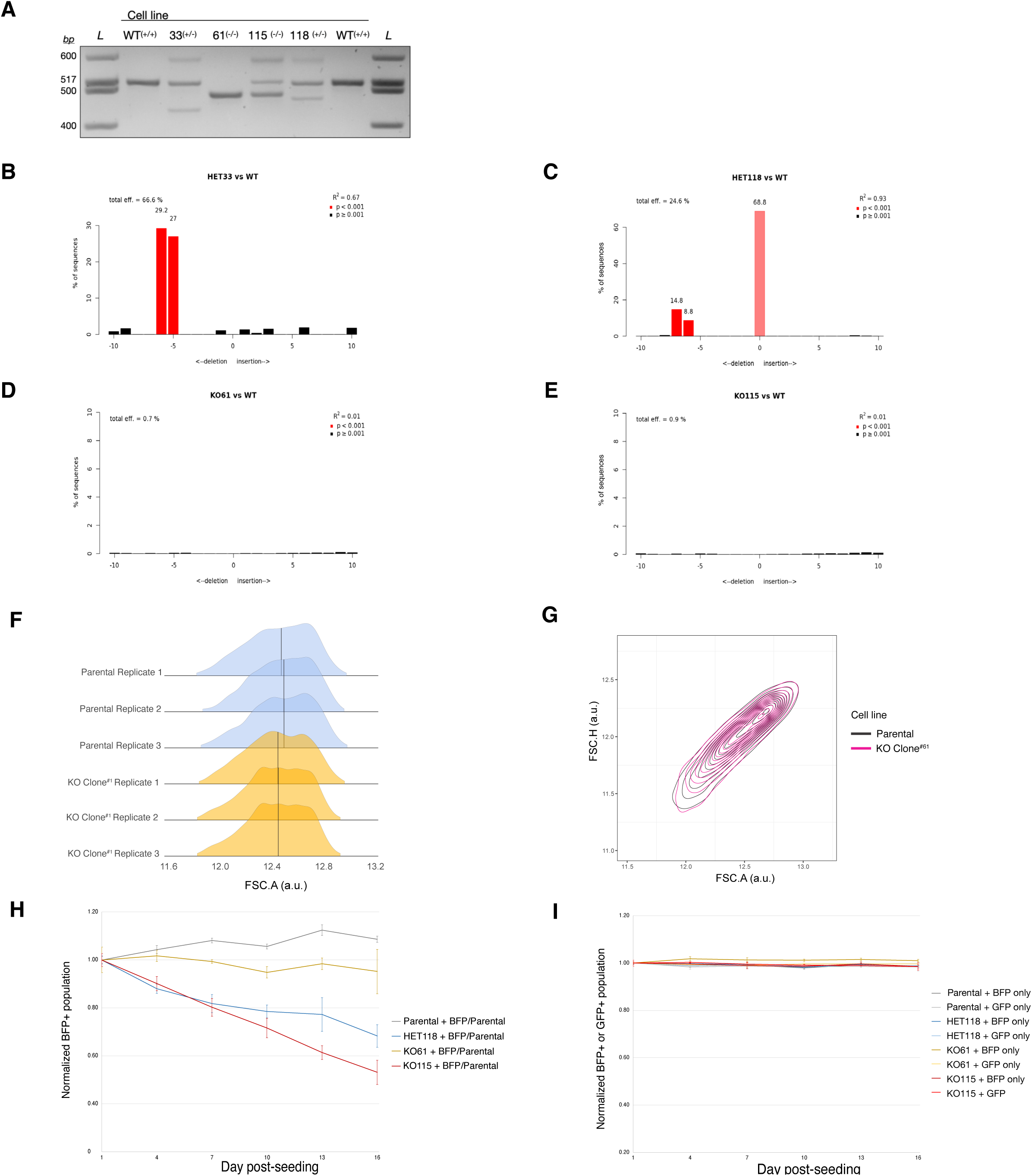
IGHMBP2 deletion clone genotyping and characterization. (*A*) Genotyping of select clones via PCR around IGHMBP2 Exon 2 cut-site. (*B-E*) TIDE alignment of Sanger sequencing results quantifying indel frequencies among alleles. (*F*) Forward-scatter area distribution between representative IGHMBP2 KO Clone#1 and parental cells in arbitrary units. (*G*) Forward-scatter area versus height profiles between representative IGHMBP2 KO Clone^#1^ and parental cells. (*H*) Competitive proliferation profiles between ΔIGHMBP2 cell lines stably expressing BFP seeded with 50% non-fluorescent parental cells. Each sample was seeded in triplicate on Day 0 and independently passaged on each day of measurement. (*I*) Measurement of BFP+ population over 2 weeks in cell lines expressing transgenic BFP or GFP. For *G* and *H*, error bars reflect the standard deviation of BFP+ populations normalized to the mean Day 1 reading among triplicate wells per sample

**Figure S2.**
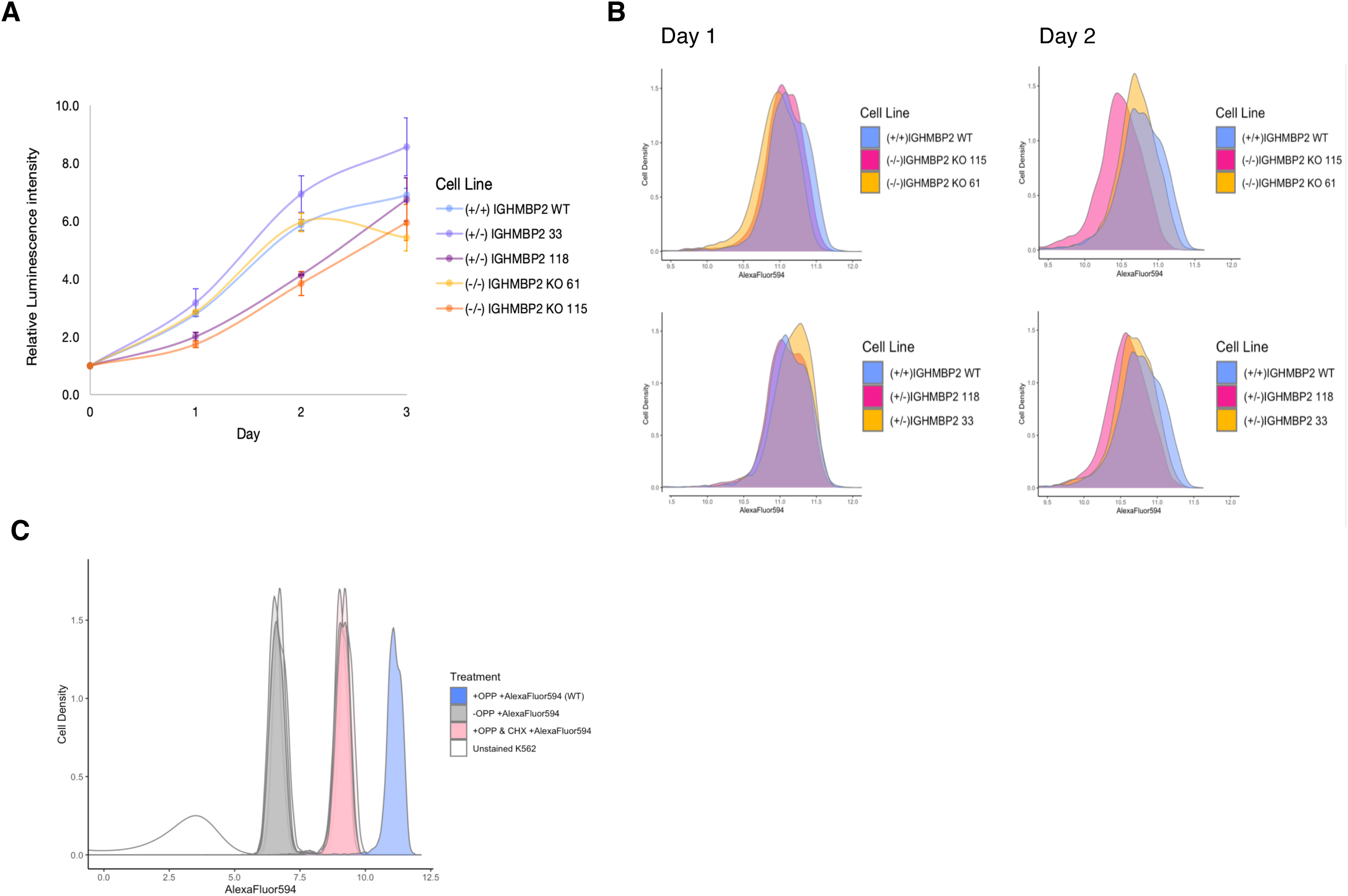
Nascent polypeptide synthesis assay with parental versus IGHMBP2 deletion clones. (A) Growth rate profiles of K562 cell lines via CellTiter-Glo metabolic assay. Error bars are standard deviation between technical replicate luminescence readings. (B) Relative levels of global translation between K562 CRISPRi parental cells versus full or partial KO of IGHMBP2 as a function of AlexaFluor594 intensities via nascent protein synthesis assay, measured by flow cytometry. Cells were harvested at growth timepoints Day 1 and 2 reflected in *A*. (C) Representative OPP assay control results showing dynamic range between unstained, cells stained with AlexaFluor594 only without OPP, cyclo-heximide-treated cells, and +OPP+AlexaFluor594 cells treated with only DMSO.

**Figure S3.**
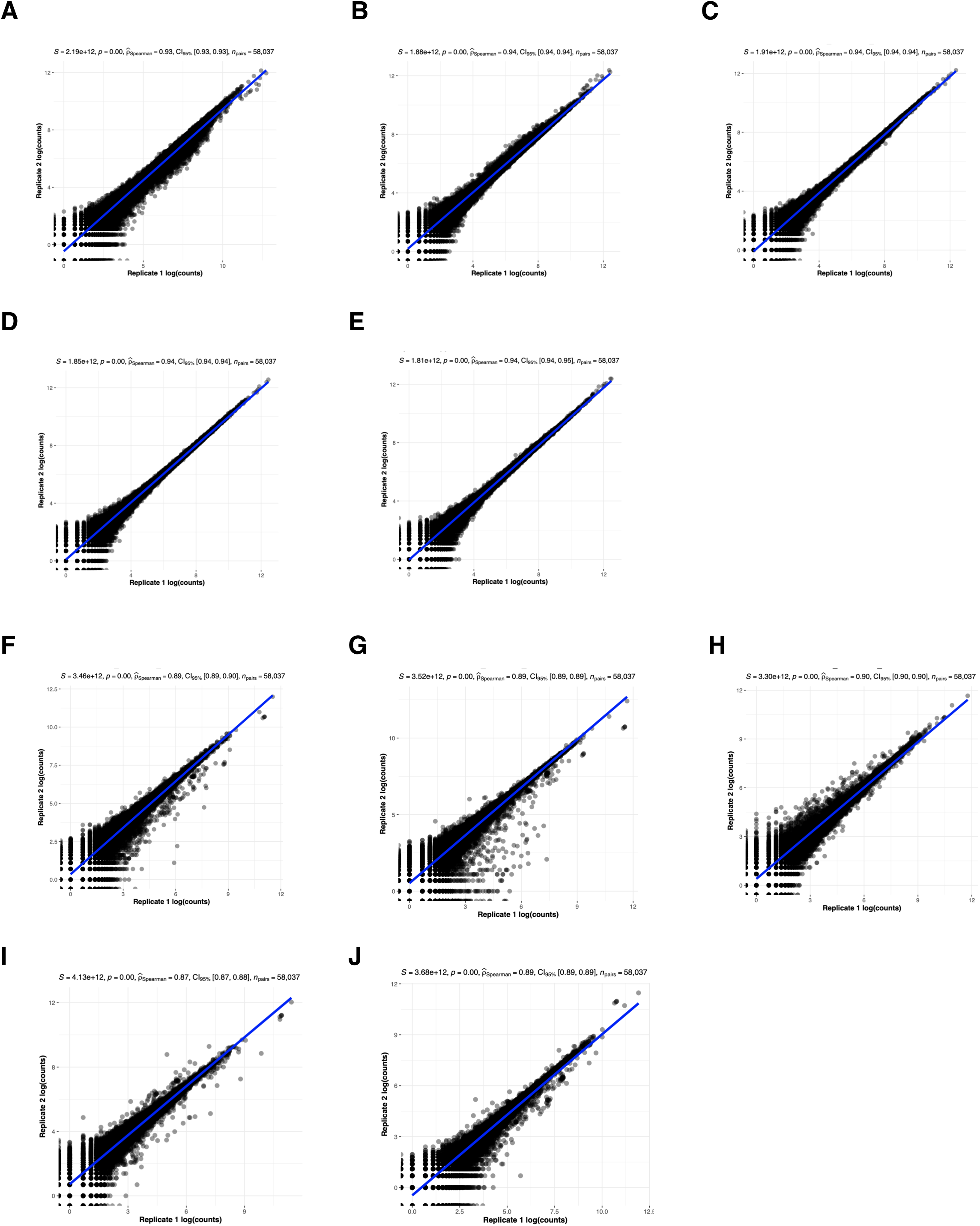
RNA-seq and Ribo-seq count correlation between replicate samples. (A) RNA-seq Spearman’s correlation analysis between parental, (B-C) HET Clones #1 and #2, and (D-E) KO Clones #1 and #2, respectively. (F) RNA-seq Spearman’s correlation analysis between parental, (G-H) HET Clones #1 and #2, and (I-J) KO Clones #1 and #2, respectively.

**Figure S4.**
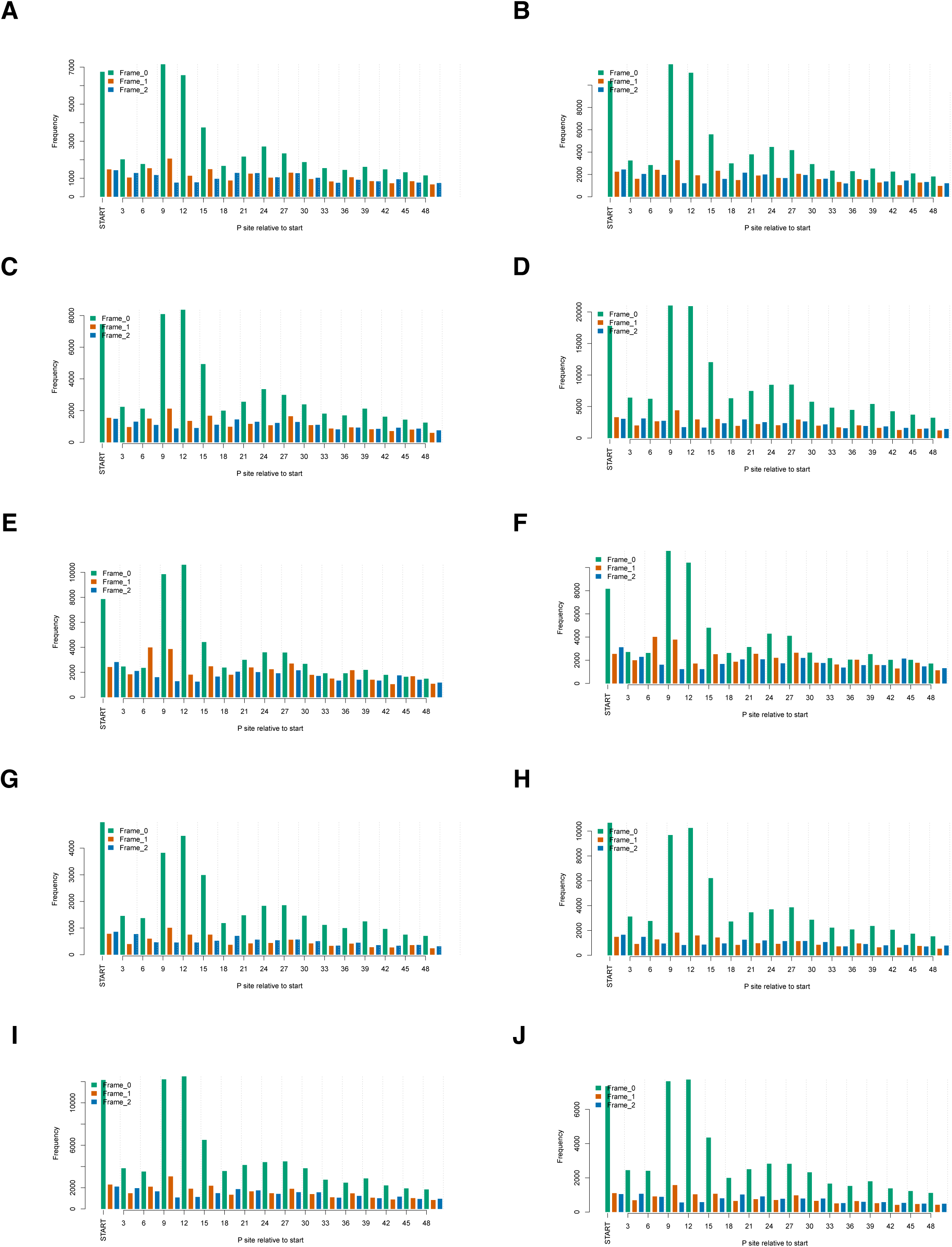
Periodicity profiles of Ribo-seq samples. (A-J) 3-nt periodicity is shown for replicates 1 and 2 of Parental, HET Clone #1, HET Clone #2, KO Clone #1, and KO Clone #2, respectively.

**Figure S5.**
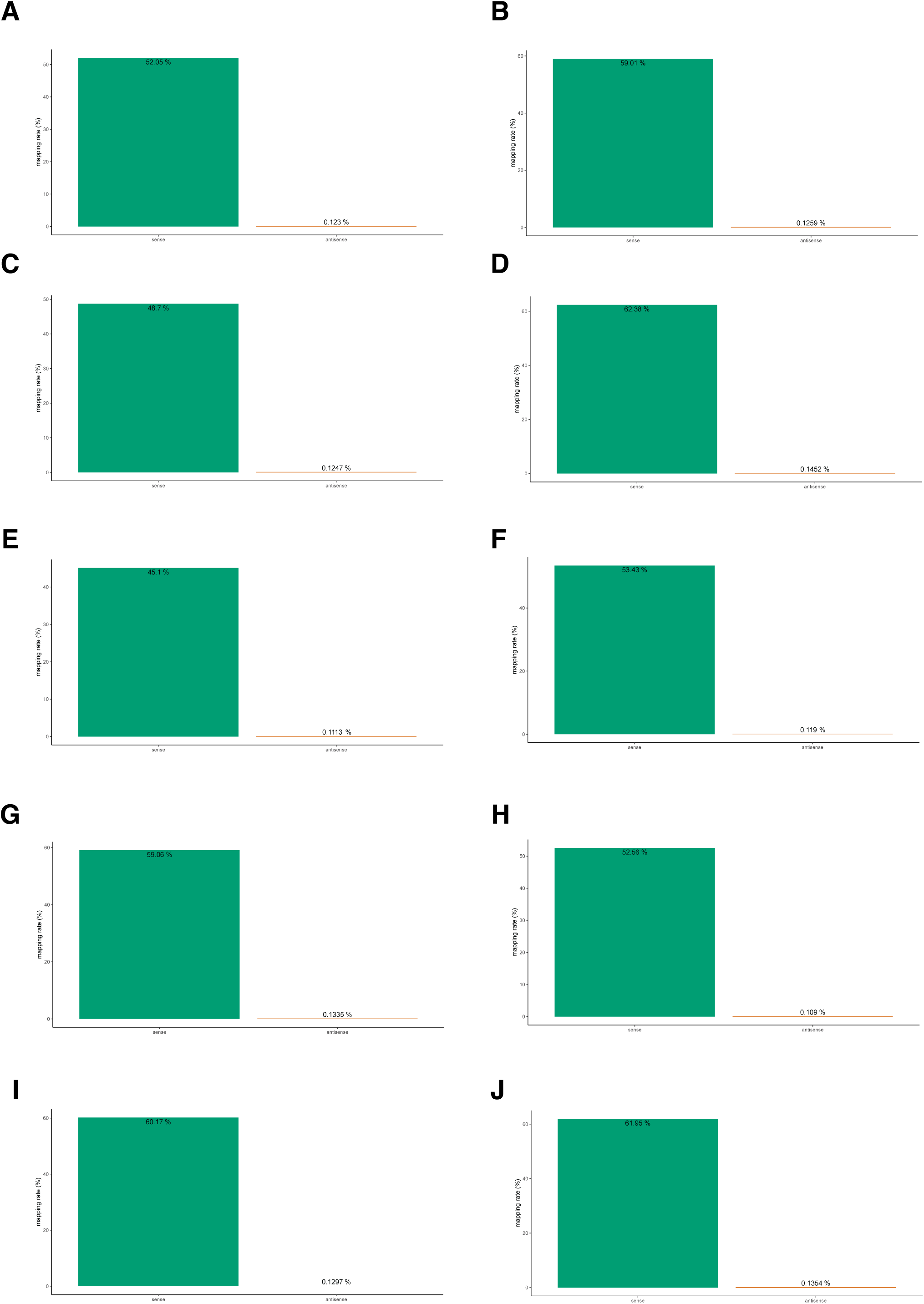
Strand sense analysis among Ribo-seq reads. (A-J) Strand sense quantification is shown for replicates 1 and 2 of Parental, HET Clone #1, HET Clone #2, KO Clone #1, and KO Clone #2, respectively.

**Figure S6.**
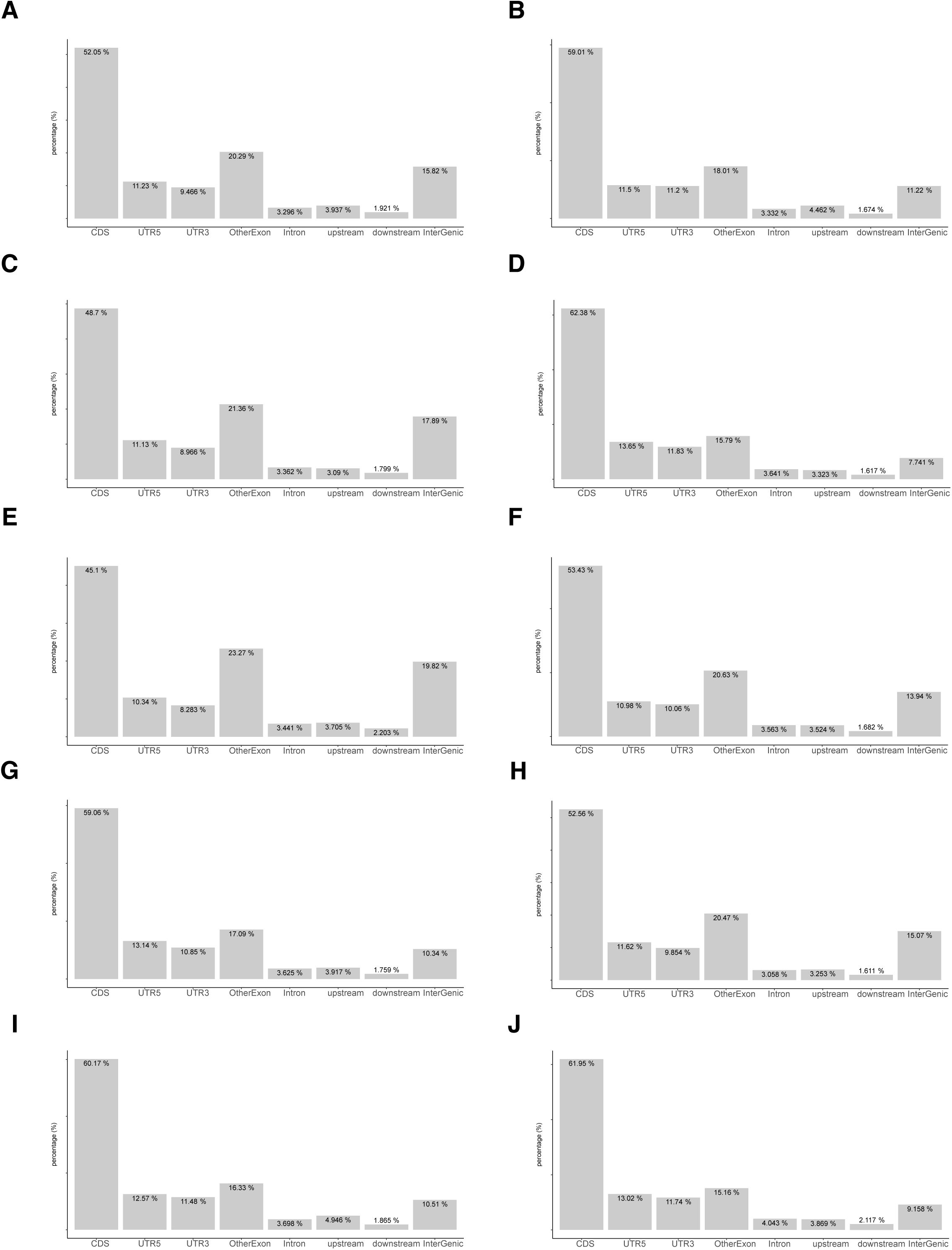
Read-mapping classifications of Ribo-seq samples. (A-J) Read distributions among CDS, UTR, and other genomic classifications shown for replicates 1 and 2 of Parental, HET Clone #1, HET Clone #2, KO Clone #1, and KO Clone #2, respectively.

**Figure S7.**
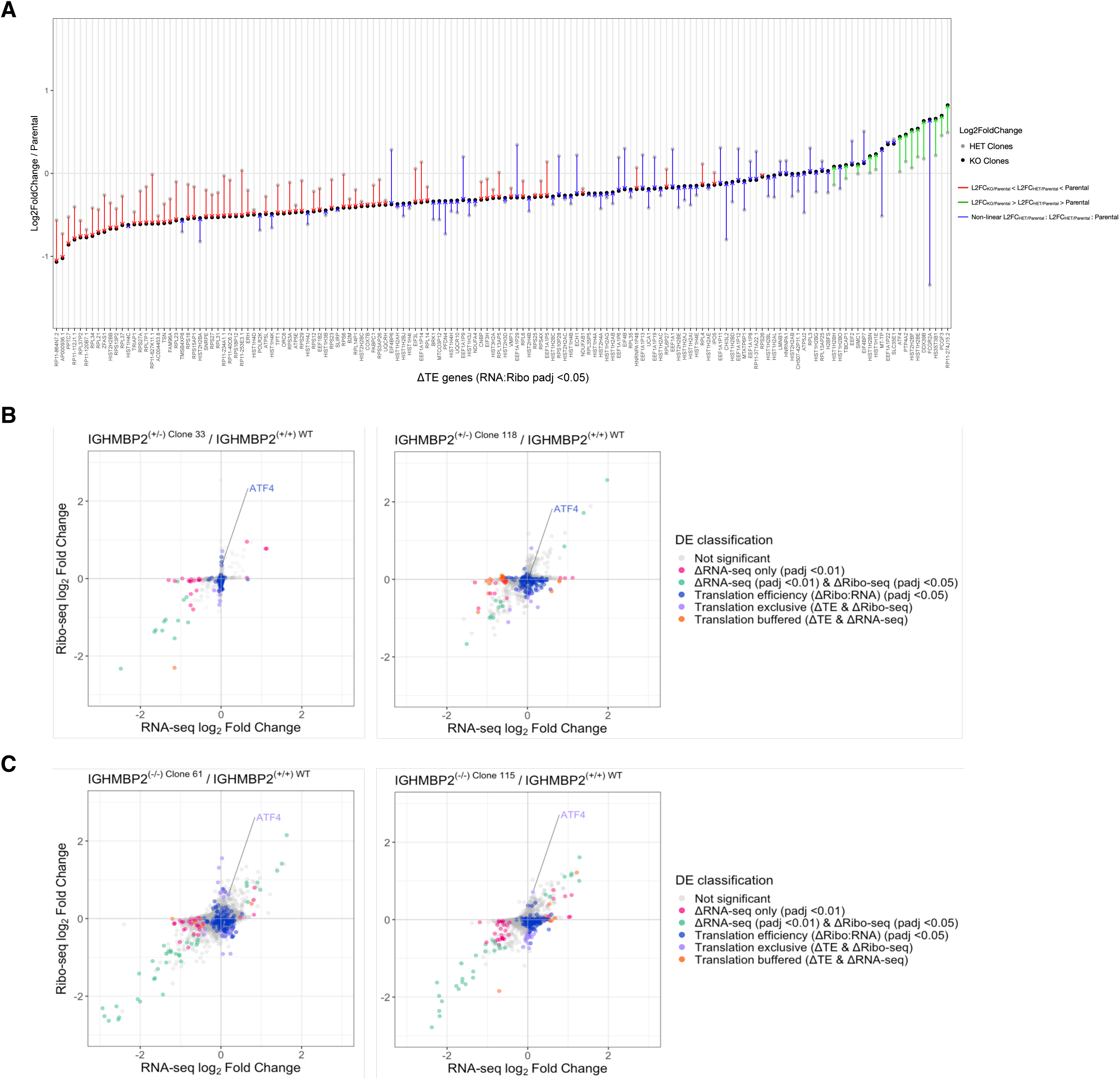
Translational efficiency of ATF4 is differentially upregulated in IGHMBP2 deletion cells. (A) Average Log2foldchanges of TE DEGs in heterozygous and full deletion clones relative to parental cells. DEGs are sorted low to high average L2FC among KO clones. Average L2FC among HET clones were then plotted and connected with lines colored by relative directionality between KO, HET, and parental conditions, where linear changes are colored red or green for down and up-regulation with respect to parental, or blue for non-linear scenarios. (B) Ribosome profiling versus RNA-seq-derived shrunken log2 fold-changes per gene in clones with partial or (C) full IGHMBP2 deletion compared to parental cells. *ATF4* is highlighted, demonstrating translation-exclusive classification is gained in IGHMBP2 KO clones with reproducible Ribo:RNA-seq positioning trends. Differential expression (DE) analysis was performed using Wald test, and p-values were adjusted (p.adj) via Benjamini-Hochberg method. Cut-offs used for DE classifications are p.adj <0.01 for Li RNA-seq (pink, green, and orange), and p.adj <0.05 for LiRibo-seq (green & violet) and Litranslation efficiency (TE; blue, violet, and orange). Genes with LiTE across all clones were identified via likelihood ratio test against the Ribo:RNA-seq interaction term. Genes of both LiTE and LiRibo-seq are identified as translation exclusive (violet). Genes of LiTE and Li RNA-seq are classified as translation buffered (orange).

**Figure S8.**
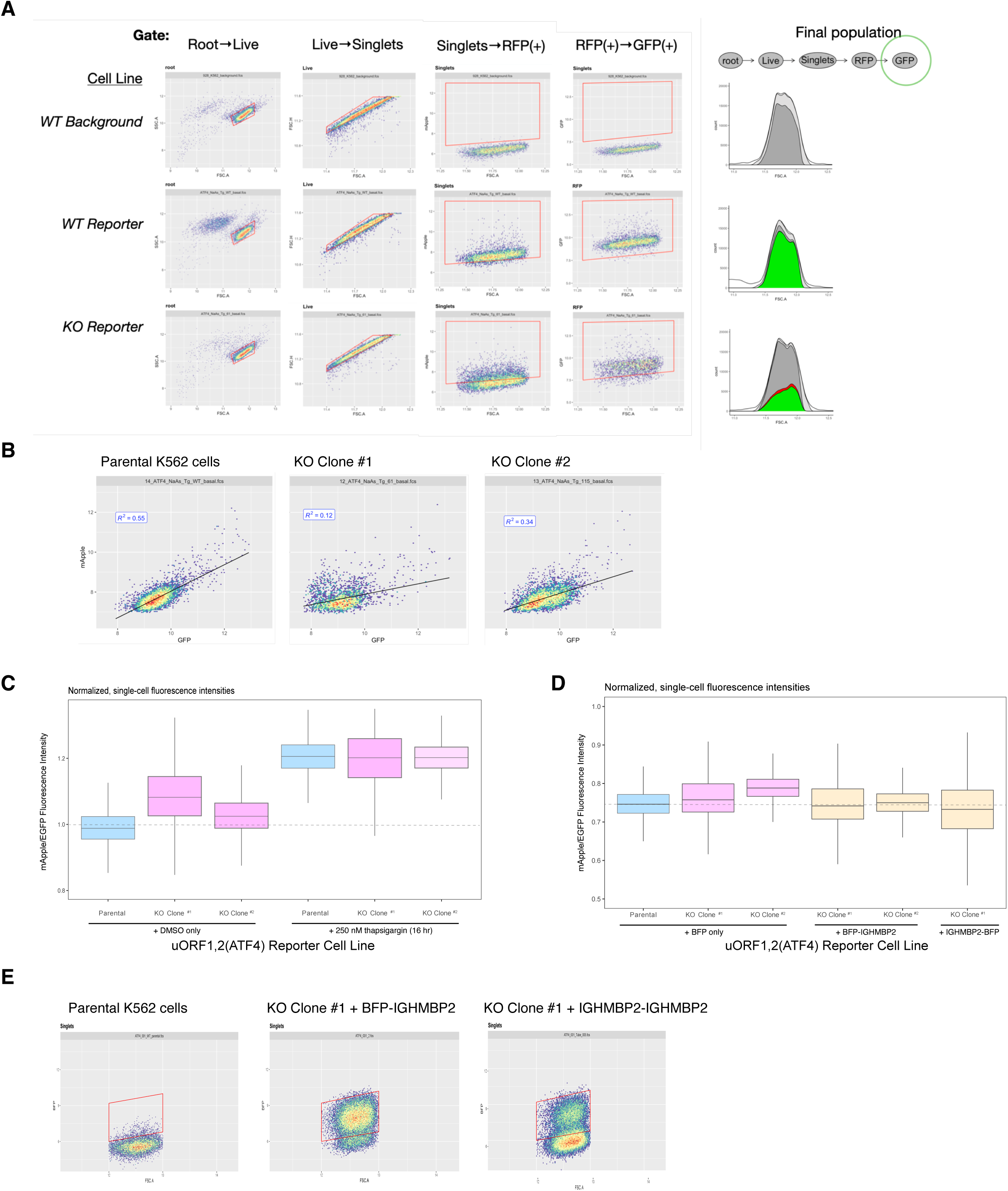
ATF4 reporter cell line characterization. (A) Gating strategy for fluorescent signals of ATF4 reporter cell lines measured via flow cytometry. (B) Correlation between GFP and mApple expression at steady-state in cell lines stably expressing ATF4 reporter. (C) uORF1,2(*ATF4*)-mApple expression normalized to promoter and translational activity (mEGFP) in LiIGHMBP2 K562 reporter cell lines at steady-state in DMSO or treated with 250 nM thapsigargin overnight. (D) Relative mApple/mEGFP intensities among reporter cell lines expressing BFP, BFP-IGHMBP2 or IGHMBP2-BFP, which were (E) gated for all BFP+ cells.

**Figure S9.**
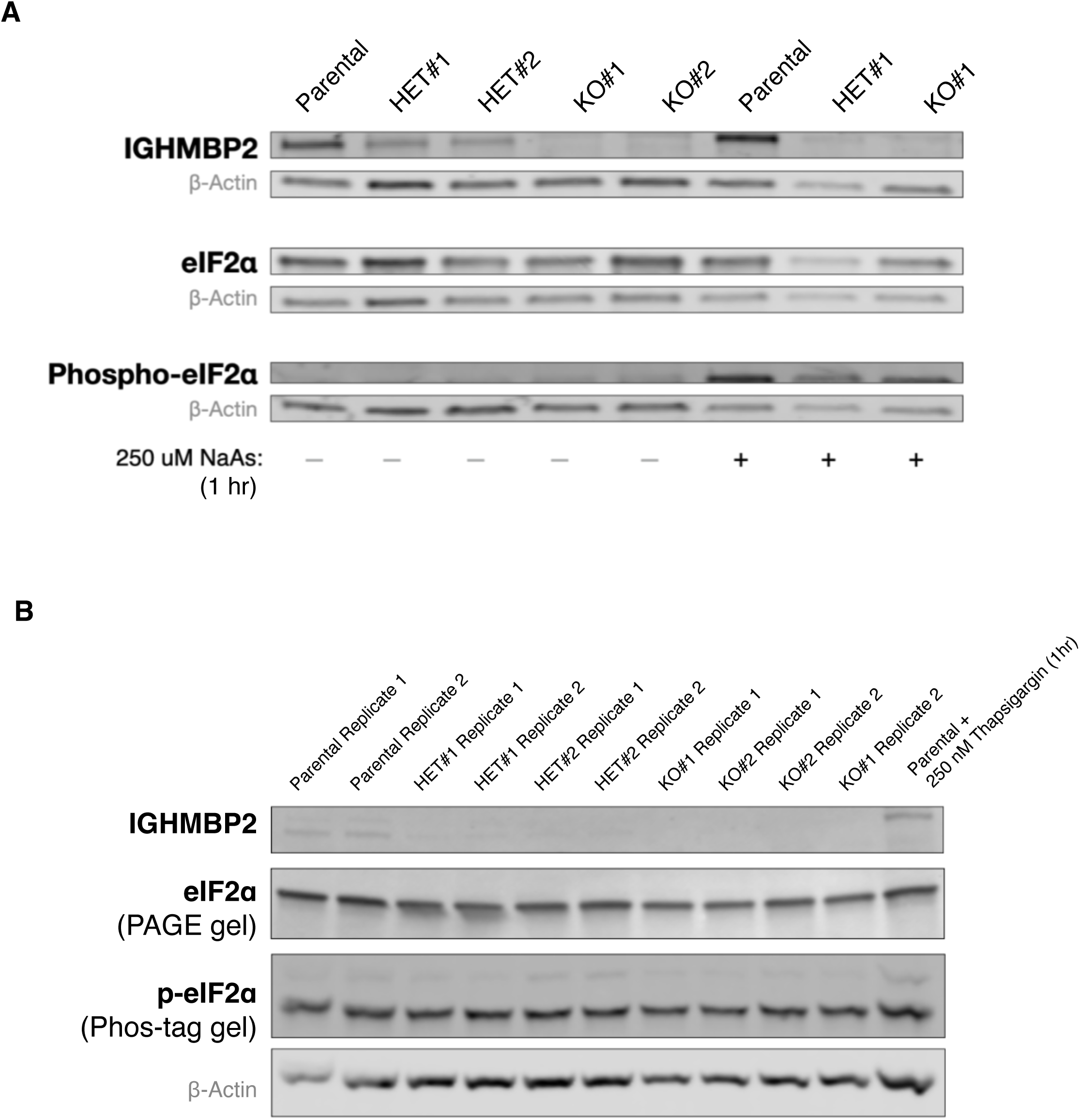
Differential p-elF2a levels at steady-state in lGHMBP2 deletion clones is not detected by Western blot. (*A*) Western blots using PAGE with 50 µg total protein loaded per lane and (*B*) Phos-Tag gels loaded with 20 µg total protein from Parental, HET Clone #1, HET Clone #2, K0 Clone #1, and K0 Clone #2 cells. *B* also shows a corresponding control PAGE gel run in parallel to confirm p-elF2a Phos-tag gel shifts are not attributed to degraded elF2a. Blots shown in *A* and *B* were incubated with anti-elF2a Ab, while a separate Western blot in A was incubated with anti-p-elF2a(S51) Ab.

**Figure S10.**
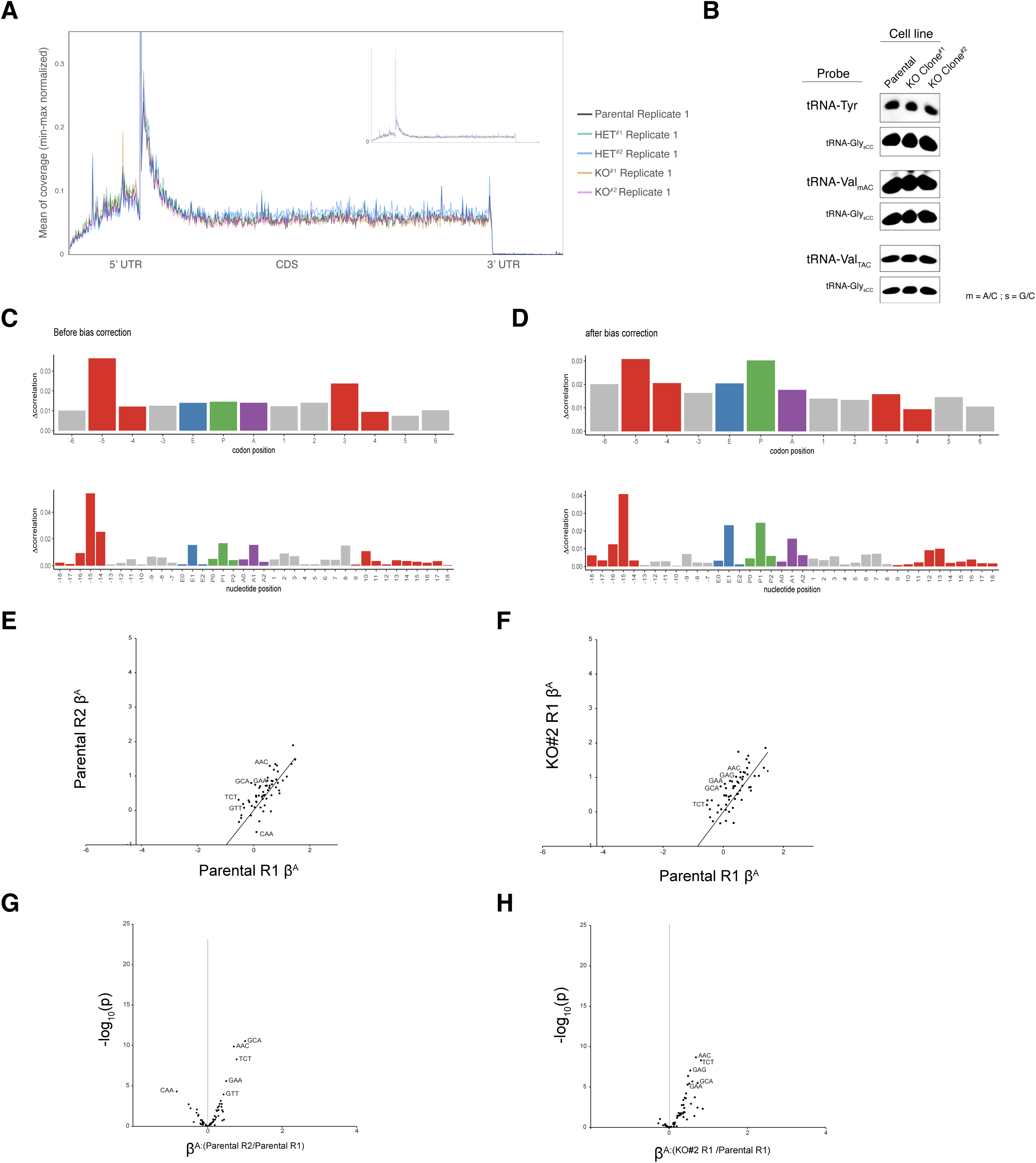
Ribosome footprint analyses across transcript regions and codons. (*A*) Representative metagene analyses with Parental, HET Clone #1, HET Clone #2, KO Clone #1, and KO Clone #2 Ribo-seq data. Zoom-out view capturing the entire TSS peak is inset. (*B*) Northern blot visualizing tRNA-Tyr (all isodecoders), tRNA-Val_mAC_ (GUU/C/G isodecoders), tRNA-Val_TAC_ (GUA isodecoders), and tRNA-Gly_sCC_ (GGC/U/G isodecoders) abundances in K562 cell lines with differential IGHMBP2 expression. 5 µg total RNA was loaded per lane. (*C-D*) Representative before and after footprint position bias-correction for Parental Ribo-seq sample processed with *choros*. (*E*) A-site codon regression coefficients between parental replicates and (*F*) KO Clone #2 relative to a parental sample. (*G*) Differential A-site codon enrichment visualized by regression coefficient interaction terms between parental replicates and (*H*) KO Clone #2 relative to a parental sample. *E* and *G* are shown to visualize expected noise among reference comparisons.

**Figure S11.**
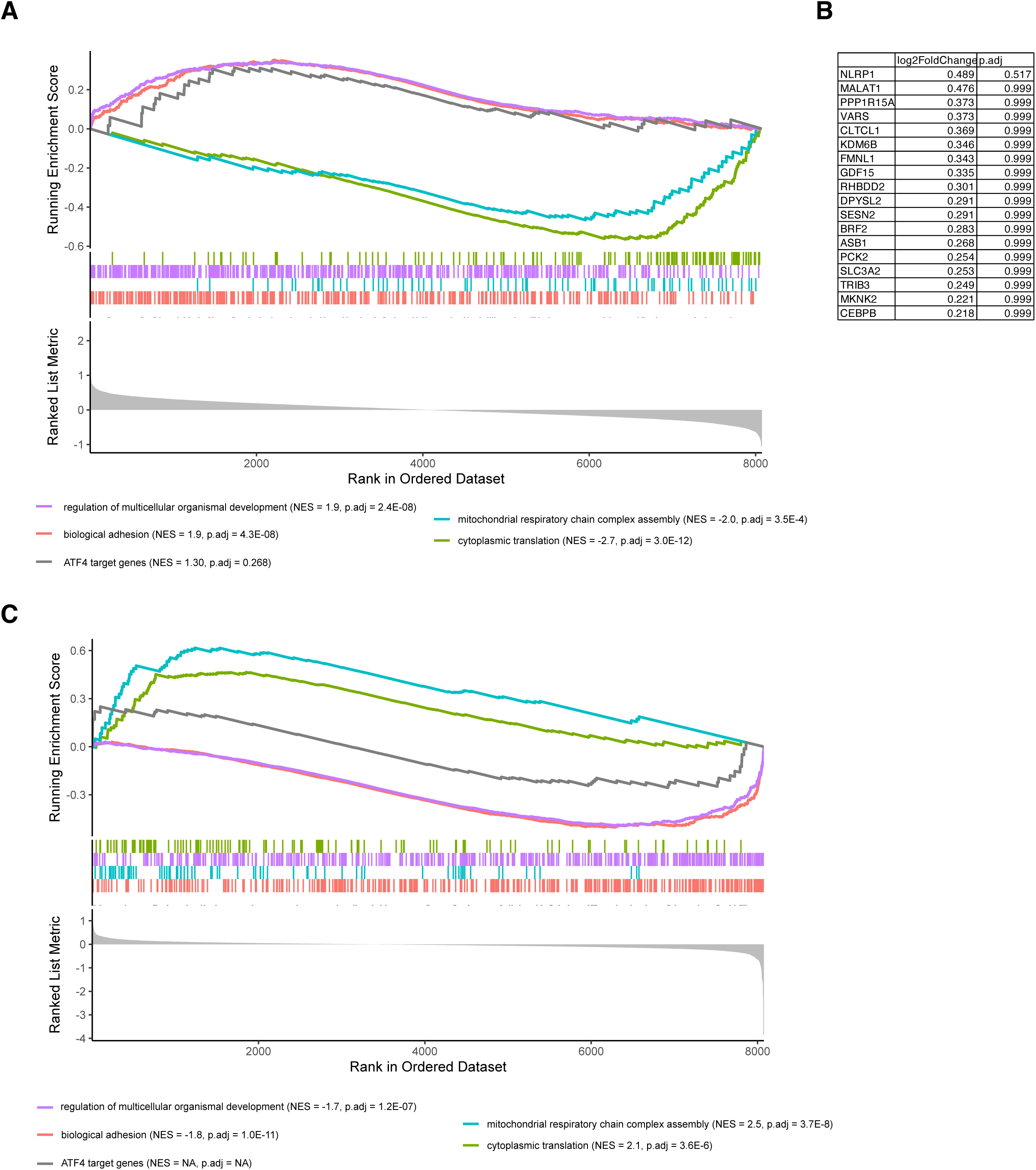
GSEA with ATF4 target gene list. (A) Rank-gene profile of ATF4 target gene list among TE DEGs from IGHMBP2 KO clones, with representative significant gene sets shown for comparison. (B) Gene set list result from ATF4 target GSE in TE DEGs from IGHMBP2 KO clones, with average L2FC between KO clones #1 and #2 and p-adjusted values per gene shown. (C) Rank-gene profile of ATF4 target gene list among RNA-seq DEGs from IGHMBP2 KO clones, with representative significant gene sets shown for comparison.GSEA was computed using the Biological Processes ontology; min GS size = 25, max GS size = 1,000 with 100,000 permutations.

